# Prospective ICH Q2(R2)-aligned total-error validation of label-free untargeted proteomics for host cell protein quantification in biotherapeutics

**DOI:** 10.64898/2026.03.06.710150

**Authors:** Somar Khalil, Jean-François Dierick, Pascal Bourguignon, Michel Plisnier

**Author notes:** Corresponding Author: Somar Khalil, Rue de l’Institut, 89, Rixensart, 1330, Belgium.

## Abstract

Untargeted proteomics enables quantitative determination of host cell proteins (HCPs) in biotherapeutics, yet no workflow has been validated under ICH Q2(R2) for regulated quality control. We report a prospective validation of label-free untargeted proteomics for HCP quantification using a total-error (TE) approach. A stable isotope-labeled whole-proteome standard was spiked into NISTmAb at seven levels (20–80 ng). Four independent assays (198 injections) supported hierarchical replication and one-way random-effects ANOVA variance decomposition with Welch–Satterthwaite adjustment. Dual entrapment analysis demonstrated empirical peptide-level false discovery proportions below 1% at q = 0.01. Deterministic parsimony inference ensured invariant protein-group definition. Weighted least-squares regression (R² = 0.993) identified stable proportional compression with recoveries of 81–85%. Repeatability dominated the variance structure (median CV 2.7%); intermediate precision total SD ranged from 0.69% to 3.81% over the validated range. Accuracy profiles integrating empirical bias with a log– log variance model showed 95% β-expectation and 95/95 content tolerance intervals fully contained within ±30%, with a lower limit of quantification (LLOQ) of 20 ng. Abundance-stratified TE analysis revealed concentration-dependent calibration heterogeneity masked by aggregate-level estimation; stratum-specific β-expectation intervals within ±35% defined an abundance-aware LLOQ of 3.6 ppm (P95 = 3.87 ppm). Robustness under independent search software (FragPipe, CCC = 0.998, LoA ±9%) and cross-platform acquisition (Astral, CCC = 0.980, LoA ±18%) remained within predefined ±30% agreement limits. System suitability criteria were derived empirically from validation performance. This is the first prospective ICH Q2(R2)-aligned validation of untargeted proteomics for HCP quantification, with a statistical framework applicable to other high-dimensional analytical methods requiring regulatory qualification.

**Graphical Abstract:** 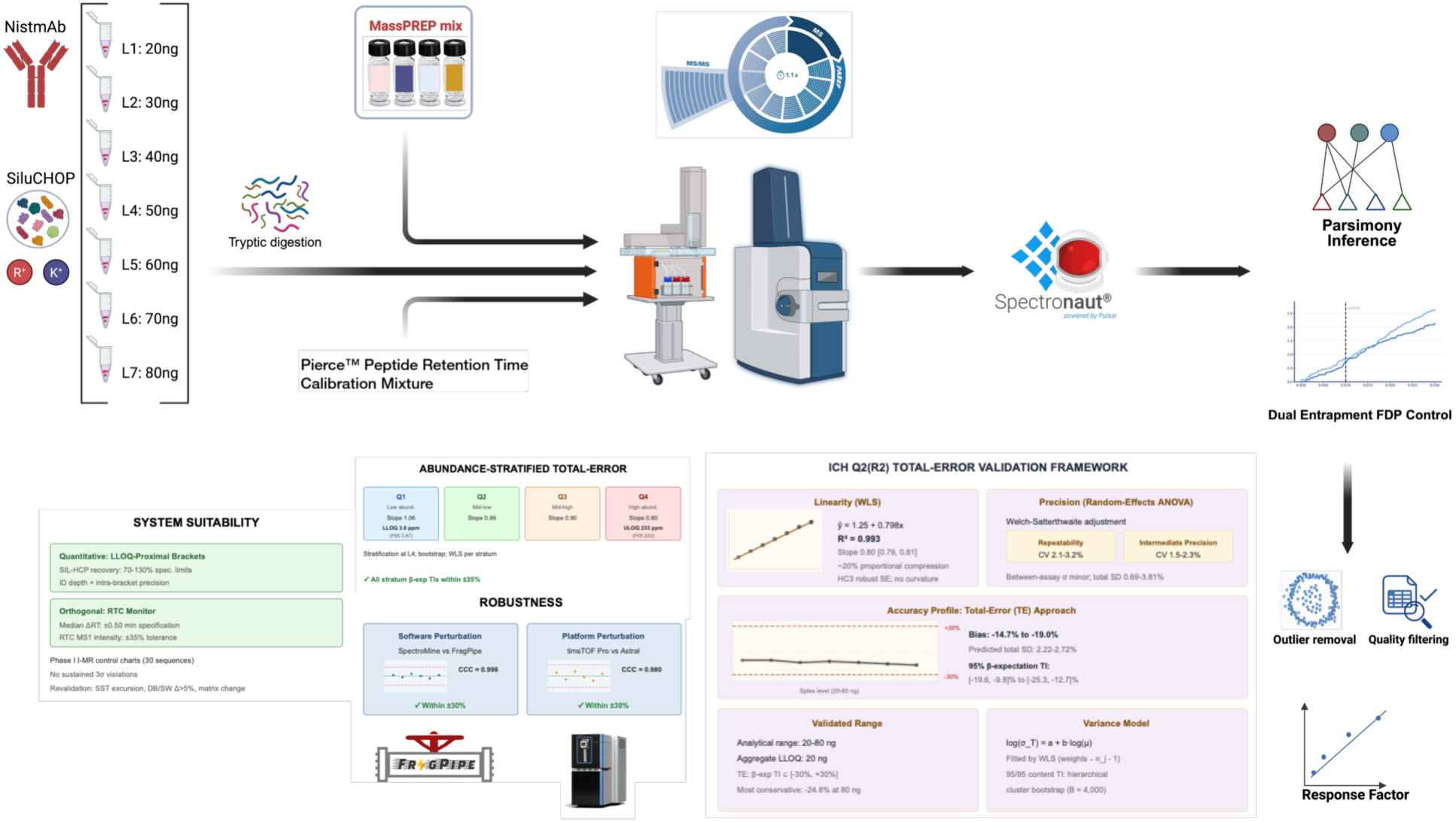

## 1. Introduction

Residual host cell proteins (HCPs) are a heterogeneous class of process-related impurities characterized by broad abundance distributions and protein-specific clearance kinetics during purification. Clinically relevant species with enzymatic or immunogenic activity may contribute minimally to total mass yet disproportionately influence risk^1–4^. Polyclonal enzyme-linked immunosorbent assays (ELISAs) generate antibody-dependent composite signals with incomplete and variable proteome coverage. A single ELISA readout is consequently poorly suited for mechanistic interpretation at the individual protein level, and orthogonal methods are now expected for complex biologics^5^.

Liquid chromatography–tandem mass spectrometry (LC–MS/MS) enables direct identification and quantification of individual HCPs^6–9^. In purified monoclonal antibody (mAb) matrices, optimized digestion and multidimensional separations can reach detection limits in the low ppm range^10–15^. The integration of trapped ion mobility with parallel accumulation– serial fragmentation (TIMS-PASEF) increases precursor sampling density and mitigates co-elution in the gas phase, improving both sensitivity and selectivity for low-abundance impurities in product-dominated backgrounds^16–22^. Within such workflows, label-free quantification based on top-peptide summation allows estimation of total HCP burden without isotopic internal standards, with quantitative performance driven by peptide selection, ionization behavior, and data-processing algorithms^23–27^.

Despite these capabilities, untargeted HCP proteomics lacks precedent as a formally validated quality control release assay under ICH Q2(R2). Label-free data-dependent acquisition (DDA) introduces abundance-dependent precursor sampling stochasticity; protein-level estimates aggregate peptides with non-random missingness; and shared sequences create inference ambiguity governed by protein-grouping rules. Database search configuration and false discovery rate (FDR) thresholds modify identification depth and downstream quantitative stability in ways not addressed by classical univariate validation constructs^28–30^. These features define a high-dimensional measurement system with a complex uncertainty structure that differs fundamentally from that of single-analyte assays traditionally addressed by ICH guidance.

ICH Q2(R2) defines validation characteristics including trueness (bias), precision (variance), linearity, and quantitation limit over a specified reporting range, and permits combined evaluation to support reliable routine performance^31^. Application to untargeted proteomics requires formal modeling of identification error and hierarchical variance components covering sample preparation, assay, analyst, and instrument platform. ICH Q14 further emphasizes lifecycle management, predefined acceptance criteria, and ongoing performance verification, requiring alignment between validation design and operational control^32^. A partial regulatory analogue exists in new peak detection (NPD) within multi-attribute methods, where untargeted signals are qualified as limit tests using predefined suitability criteria and spike-based detection limits^33^. NPD, however, operates on a fixed product digest with an established reference profile and does not generalize to HCP proteomics, where the set of reportable proteins is database-derived and inference-driven.

Current USP ⟨1132.1⟩ provides operational guidance for MS-based HCP analysis and describes three quantitative strategies (relative to product protein, spiked intact proteins, and stable isotope-labeled peptides)^34^. The chapter distinguishes product-specific ICH-aligned method validation from a broader system qualification approach but does not define the measurand for untargeted outputs in metrological terms, nor establish a statistical framework linking database composition, FDR control, protein grouping, and system suitability criteria to validated performance characteristics. Quantitation limits are described using ICH Q2(R1)-aligned terminology and are not extended to integrated bias–variance evaluation under Q2(R2). USP ⟨1132.1⟩ therefore establishes technical best practices for MS-based HCP analysis but does not articulate a validation architecture for inference-conditioned proteomic measurement systems under ICH Q2(R2).

Recent work has demonstrated that the three USP ⟨1132.1⟩ quantitative strategies can be implemented within an ICH Q2(R2)-aligned qualification design for a small number of predefined HCPs spiked into a mAb matrix^35^. In that design, the measurands (protein identity, surrogate peptides, and concentration levels) are specified *ex ante*, and validation characteristics are estimated for these fixed analytes within a controlled configuration. These results confirm that MS-based HCP assays can satisfy Q2(R2) performance criteria for targeted proteins. Such designs do not, however, address the inference-dependent analyte space of untargeted HCP proteomics, where protein identities arise from database search and grouping rules. Detectability, grouping stability, and quantitative behavior within the endogenous HCP population remain uncharacterized, and extrapolation from two spike-defined proteins to hundreds of database-defined species presumes quantitative homogeneity that has not been empirically demonstrated.

For untargeted HCP workflows, validation must operate at the level of the measurement system. In practice, this means the validation must demonstrate that identification error is controlled empirically, that performance is characterized across the abundance range, and that protein groupings remain stable when database and inference parameters are fixed. Within an ICH Q2(R2)/Q14 framework, system-level validation integrates total-error (TE)–based accuracy profiling, predictive intervals with prespecified coverage probabilities, and hierarchical variance-component estimation to define the reportable range, lower limit of quantitation (LLOQ), and system suitability criteria linked to continued performance verification.

We present a prospective, hierarchical validation of label-free ddaPASEF proteomics for quantitative determination of HCPs in mAb matrices, aligned with ICH Q2(R2) and implemented within the TE approach^36^. The design uses an absolute nanogram load scale and a stable isotope-labeled whole-proteome standard as a compositionally representative calibrant^37^. Three interlocking components support the TE framework: empirical FDP control via dual entrapment, deterministic parsimony for stable protein-group assignment, and random-effects ANOVA with Welch–Satterthwaite adjustment. These are integrated to derive β-expectation and 95/95 content tolerance intervals (TIs) that combine trueness and precision into unified acceptance criteria.

Abundance-stratified TE profiling defines an abundance-aware validated range and LLOQs. Cross-software and cross-platform comparisons evaluate robustness of the integrated analytical–computational system^38^. Empirically derived system suitability tests (SSTs) and longitudinal process control connect study-phase performance to routine monitoring. To our knowledge, this is the first prospective ICH Q2(R2)-aligned validation of untargeted proteomics for HCP quantification. The statistical framework is transferable to other high-dimensional analytical methods requiring regulatory qualification.

## 2. Experimental Section

### 2.1 Validation Design

Validation employed a prospective hierarchical design aligned with ICH Q2(R2) for quantitative impurity assays and adapted to a high-dimensional LC–MS/MS measurement system comprising sample preparation, acquisition, database search, protein inference, and quantitative summarization (**Figure 1**; **Table 1**)^31,36,39^.

**Figure 1:**
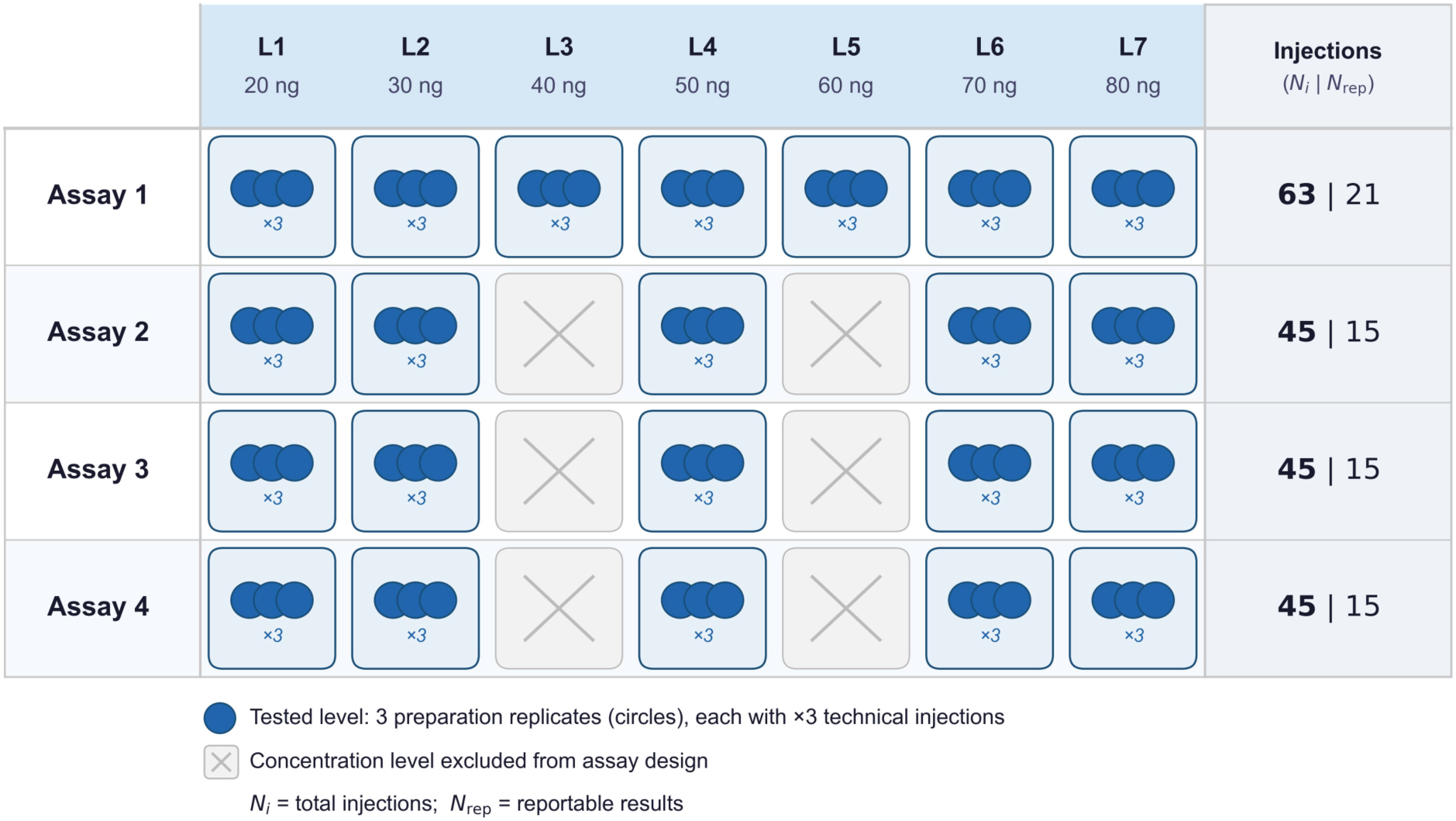
Hierarchical validation design for label-free ddaPASEF HCP quantification. Seven spike levels (20–80 ng) were evaluated across four independent assays. At each included level, three independent preparations were analyzed in technical triplicate (3 × 3). Grey tiles indicate levels not included in a given assay. The right column reports total injections (Nᵢ) and replicate-block reportable results (N₍rep₎), each defined as the arithmetic mean of three technical injections.

**Table 1.** Sources of analytical variation incorporated into the validation and robustness design.

| Factor | Levels | Role |
| --- | --- | --- |
| Assay run | 4 independent assays | Between-assay variance |
| Analyst | 3 analysts | Operational robustness (absorbed within-assay variance) |
| Independent preparation | 3 per level per assay | Sample preparation variance |
| Technical injections | 3 per independent preparation | Instrument repeatability |
| LC column | 3 columns | Chromatographic robustness |
| LC–MS system (validated system) | Evosep One + timsTOF Pro | Primary validated measurement system |
| LC–MS system (robustness comparison) | Vanquish Neo + Orbitrap Astral<sup>38</sup> | Cross-platform robustness |
| Software pipeline (validated system) | SpectroMine | Primary processing workflow |
| Software pipeline (robustness comparison) | FragPipe | Algorithmic robustness |

The validated measurand is the aggregate total HCP mass estimated by Hi3 label-free quantification and reported as nanograms of HCP per injection under the defined SIL-HCP calibration and deterministic protein-group inference rules. The measurement system generating this quantity comprised sample preparation, LC–MS/MS acquisition (Evosep One– timsTOF Pro operated in ddaPASEF mode), database search with FDR control, deterministic protein inference, Hi3-based quantification, and post-processing required to compute the reportable total HCP value. All analytical and computational parameters were version-locked prior to study initiation.

The design was defined *a priori* to support variance decomposition and TE estimation at the replicate-block level. Per spike level, replicate-block relative error was modeled using a one-way random-effects model with assay run as the grouping factor to estimate between-assay and within-assay components. The within-assay component aggregates variation from preparation, analyst, LC column, and residual effects under the locked configuration. Four independent assay runs were performed. Assay 1 included a complete seven-level spike series (20–80 ng). Assays 2–4 included five levels (L1, L2, L4, L6, L7). At each level, three independent preparations were analyzed in technical triplicate (3 × 3). The arithmetic mean of each triplicate defined the reportable result (N₍rep₎); total injections (Nᵢ) denote all raw LC–MS/MS acquisitions. Levels present in all four assays yielded twelve replicate-block observations spanning assay runs, analysts, and LC columns.

Cross-platform robustness was evaluated by comparing total HCP values for a subset of samples generated on an Evosep One–timsTOF Pro system with those obtained on a Vanquish Neo–Orbitrap Astral platform under matched spike levels and harmonized processing parameters^38^. Software robustness was assessed by reprocessing identical raw files using an independent search engine, FragPipe.

### 2.2 Analytical procedure

The locked protocol is provided in **Supplementary Method S1**. NISTmAb (RM 8671) served as matrix. A stable isotope–labeled CHO whole-proteome standard (SIL-HCP; Sigma) was spiked at seven levels (20–80 ng total HCP injection load). Samples underwent denaturation, reduction, alkylation, and overnight digestion with trypsin/Lys-C (1:40, w/w), followed by C18 solid-phase extraction. Four MassPREP protein digest standards were added post-digestion for response-factor determination. Peptides (500 ng) were separated on an Evosep One (30 samples/day method) coupled to a timsTOF Pro operated in ddaPASEF mode. Database searching was performed in SpectroMine against the Cricetulus griseus proteome (trypsin/P specificity, ≤1 missed cleavages) with 1% FDR control at PSM, peptide, and protein levels.

Protein inference was performed using a deterministic parsimony algorithm retaining groups supported by ≥1 unique peptide; lead accessions were defined by maximal peptide evidence. The full algorithmic specification is provided in **Supplementary Method S2**; the inference script is archived at DOI: 10.5281/zenodo.18826281.

Peptide-level filtering included single-hit exclusion, modified Z-score outlier removal^40^, intensity deviation screening, total ion current normalization, and replicate CV filtering. Protein abundance was estimated using the Hi3 approach^27^. Absolute HCP mass was calculated as:

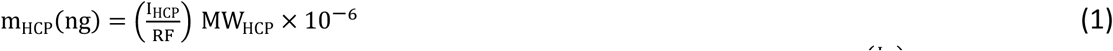

where I_HCP_ is the summed Hi3 intensity of the protein and 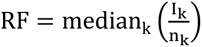 is the response factor derived from the MassPREP standards (I_k_ Hi3 intensity; n_k_ injected amount in fmol). MW_HCP_ is the molecular weight of the lead protein. Total HCP is defined as the sum of all inferred protein-group masses and constitutes the reportable value used for validation analyses^23,27,28^.

### 2.3 Empirical estimation of false discovery proportion

Peptide-level specificity was evaluated independently of Target-decoy competition using a search-embedded entrapment strategy^41^. Two orthogonal entrapment spaces were constructed. The first comprised shuffled *C. griseus* tryptic peptides generated by permuting internal residues while preserving proteolytic termini. The second comprised a trimmed foreign-proteome space derived from *in silico* digestion of Arabidopsis thaliana and additional non-CHO proteomes, with peptides overlapping the CHO peptidome removed. Each entrapment database was constructed to approximate a 1:1 ratio (r) of unique entrapment to unique target peptides. Entrapment FASTAs were concatenated with the target FASTA and searched under identical SpectroMine parameters.

PSM-level results were collapsed to unique stripped peptide sequences by retaining the minimum q-value per sequence. For a peptide-level threshold τ, the empirical FDP was estimated as:

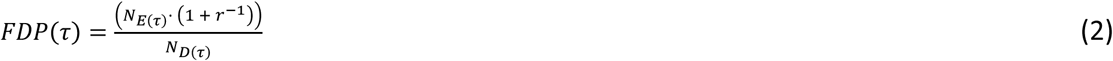

where N_E(τ)_ is the number of unique entrapment peptides at or below τ, and N_D(τ)_ is the total number of unique peptides (target + entrapment) at or below τ. For r = 1, the estimator reduces to 2N_E_(τ)/N_D_(τ).

FDP was evaluated over the reporting range of τ. Uncertainty was quantified by nonparametric bootstrap at the peptide level: stripped sequences were resampled with replacement, FDP recomputed per resample, and pointwise 95% percentile bands derived. At τ = 0.01, uncertainty in the entrapment proportion *N*_*E*_/*N*_*D*_ was additionally summarized using a two-sided 95% Wilson score interval and scaled by (1 + r^−1^).

### 2.4 Statistical analysis

The complete statistical analysis pipeline is archived at Zenodo (DOI: 10.5281/zenodo.18826281). Data processing and numerical operations used NumPy (v2.0.1) and pandas. Statistical procedures, including random-effects ANOVA, weighted least-squares (WLS) regression, hierarchical bootstrap resampling, kernel density estimation (KDE), and tolerance-interval construction, were implemented in SciPy and statsmodels. Figures were generated in Matplotlib. Bootstrap procedures employed a fixed NumPy random seed (0). Final figure layout refinement was performed in GraphPad Prism (v11.0).

#### 2.4.1 Linearity

Linearity of total HCP quantification was evaluated by regressing measured total HCP (ng) on nominal spike amount over L1–L7 (20–80 ng) using replicate-block reportable results pooled across assays. The WLS model was specified as:

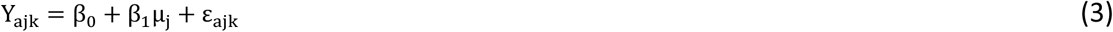

where Y_ajk_ denotes the replicate-block mean at assay a, spike level j, and replicate block k, and µ_j_ is the nominal spike amount. Weights were defined as:

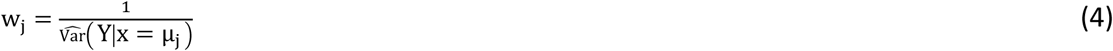

where 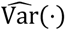 is the empirical variance of replicate-block reportables at level j. Coefficient uncertainty was estimated using HC3 heteroscedasticity-consistent standard errors^42^.

Linearity was evaluated by testing the null hypotheses *H*_0_: *β*_0_ = 0 and *H*_0_: *β*_1_ = 1. Model adequacy was assessed by examination of residual patterns and variance behavior over the concentration range.

#### 2.4.2 Accuracy profile

Accuracy of total HCP was evaluated on the relative-error scale within a TE framework aligned with Société Française des Sciences et Techniques Pharmaceutiques (SFSTP) guidance^36,39^. Reportable results were defined at the replicate-block level as the mean total HCP (ng) across technical injections within each assay and spike level. Relative error was defined as:

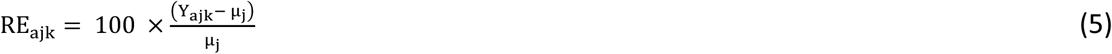

where Y_{ajk}_ is the replicate-block mean and µ_j_the nominal spike amount at level j. Level-specific bias was estimated as:

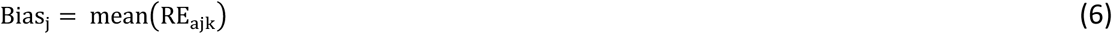

Bias was modeled as a level-wise step function without parametric smoothing.

##### 2.4.2.1 Variance decomposition and modeling

At each spike level, variance on the relative-error scale was estimated using one-way random-effects ANOVA with assay as grouping factor^39^. The within-assay variance component 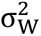 (repeatability) was taken as MS_within_. The between-assay variance component 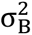 was estimated by method of moments using harmonic-mean replication m_h_:

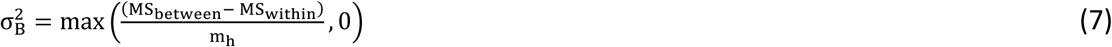

Total SD (intermediate precision under varied conditions) was:

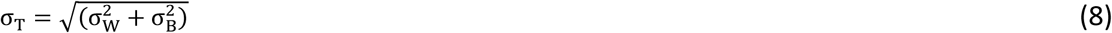

Level-specific total SDs were modeled by a log–log relationship:

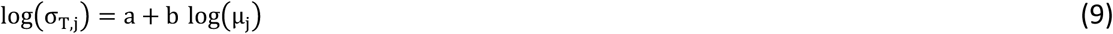

fitted by weighted least squares with weights proportional to n_j_ − 1, where n_j_is the number of replicate-block observations at level j. Predicted SDs 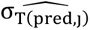 were used for TI construction.

##### 2.4.2.2 Tolerance interval construction and acceptance criteria

The TE approach evaluates whether future routine measurements fall within predefined acceptability limits by integrating systematic bias and random variability into a single decision criterion. At each spike level, two TIs were estimated. The 95% β-expectation interval describes the range within which a future replicate-block relative error is expected to fall with probability β given the observed bias and variance structure. In parallel, a 95/95 content TI was estimated to provide 95% confidence that at least 95% of future errors are contained within the reported bounds. A level was considered validated when the β-expectation interval was fully contained within the predefined ±30% acceptance limits.

For each level:

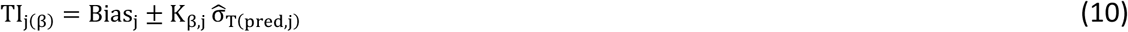

With β = 0.95. The coverage factor K_β,j_ was derived from the Student t distribution using Welch– Satterthwaite effective degrees of freedom applied to the ANOVA mean-square representation of the total variance estimator:^43^

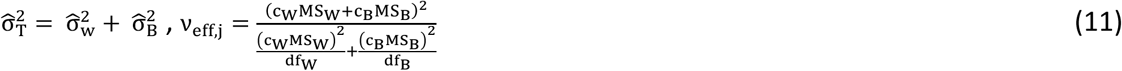

where c_W_ = 1 − 1/m_h_, c_B_ = 1/m_h_, df_W_ = N_j_ − A_j_, df_B_ = A_j_ − 1, A_j_ is the number of assays and N_j_ the number of replicate-block observations. Effective degrees of freedom were bounded below by 3. The coverage factor was:

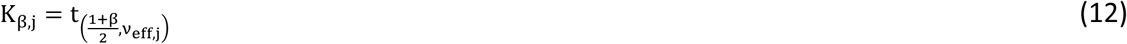

In parallel, 95/95 content TIs were estimated using a hierarchical cluster bootstrap with parametric simulation of future errors^44^. For B = 4000 iterations, assays were resampled with replacement; replicate blocks were resampled within assay at each level. Bias and variance models were refitted per bootstrap sample. Simulated future errors were generated as:

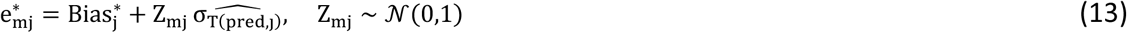

At each level, the inner half-width was defined as the 95th percentile of 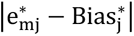 over simulated values. The 95/95 half-width HW_j(95/95)_was the 95th percentile of these inner half-widths across bootstrap replicates. The final interval was:

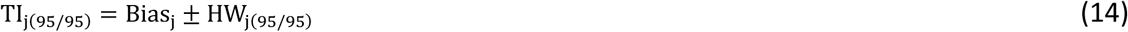

By construction,

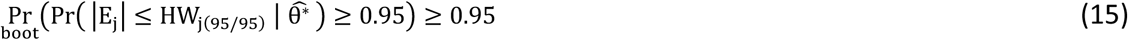

where E_j_ denotes a future replicate-block relative error and 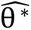 the bootstrap-refitted bias and variance parameters.

A level was considered validated when:

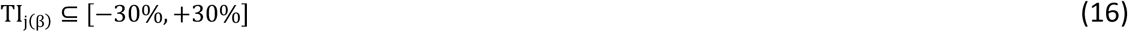

The ±30% aggregate acceptance limit was derived from prior performance characterization and reflects the combined uncertainty expected for a label-free untargeted LC–MS/MS workflow operating on a multi-protein calibrant across independent assay runs (**Supplementary Note S4**). This limit accommodates known sources of systematic compression in Hi3 quantification, including response-factor dispersion across the SIL-HCP proteome, incomplete digestion equivalence, and ionization-dependent biases, while preserving analytical relevance for aggregate HCP burden estimation^27,45^. The wider ±35% limit applied to abundance-stratified analysis accounts for the additional variance introduced by bootstrap-based stratum-level estimation and the smaller effective protein populations per stratum. Both limits were predefined prior to data analysis and align with TE acceptance criteria commonly applied in quantitative bioanalytical validation.

#### 2.4.3 Abundance-stratified total-error analysis

Protein abundances were expressed in ppm (ng HCP per mg mAb). Stratification was defined once using the L4 spike level. For each protein p, the L4 reference abundance 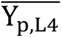 was computed as the mean ppm at L4 pooled over assays and technical replicates. Fixed strata (Q1– Q4) were assigned using empirical quantiles of 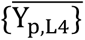 with cutpoints at the 5th, 25th, 50th, 75th, and 100th percentiles. The 5th-percentile lower bound limited leverage from extremely low-abundance proteins with unstable estimates. Stratum membership was fixed for all subsequent analyses.

For each assay a, spike level j, replicate block r, and stratum b, a stratum-level reportable was obtained by nonparametric bootstrap over proteins within cell (a, j, r, b). Let 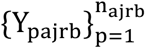 denote ppm values in that cell. For B bootstrap iterations, proteins were resampled with replacement and the mean computed. The reportable was the bootstrap expectation:

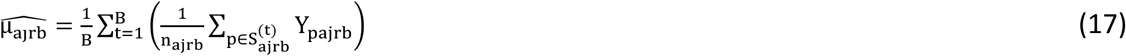

Cells with fewer than 30 proteins were excluded.

A stratum-specific normalization anchor was defined as the mean L4 reportable pooled across assays and replicate blocks:

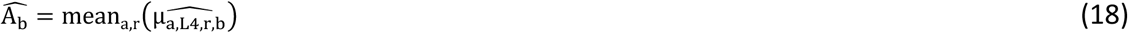

Observed and theoretical spike ratios were:

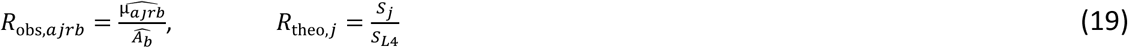

Relative error on the ratio scale was:

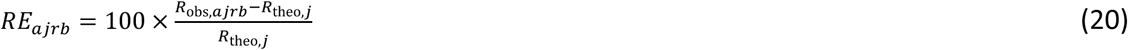

Within each stratum, TE accuracy profiles were constructed on RE_ajrb_. Level-specific bias was estimated as the empirical mean at each spike ratio. Variance was decomposed using one-way random-effects ANOVA with assay as grouping factor. Total SD was modeled by a log–log relationship in the theoretical spike ratio x_norm_ = S_j_/S_L4_.

The 95% β-expectation TI was:

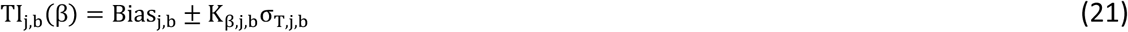

where 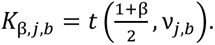

with β = 0.95 and ν_j,b_ obtained by Welch–Satterthwaite (minimum df = 3). A spike ratio was accepted within stratum b when:

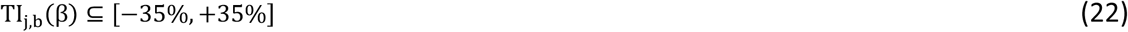

The validated domain was defined as the intersection of acceptable spike ratios across strata. Abundance-aware quantification limits were derived from validated levels. The LLOQ was defined as the 95th percentile of stratum-level reportables at the lowest validated spike ratio within Q1. The ULOQ was defined as the 5th percentile of reportables at the highest validated spike ratio within Q4.

All TI calculations were performed on dimensionless ratios; ppm units are reported for interpretability only and do not affect the statistical evaluation.

#### 2.4.4 System suitability testing

SST was incorporated into the analytical procedure to verify continued quantitative and identification performance during routine operation. SST evaluates performance at the lower boundary of the validated range and differentiates quantitative degradation from chromatographic or instrument-state perturbation.

The primary SST component comprised bracketed acquisitions of SIL-HCP at the LLOQ-proximal level (L1; 20 ng total HCP). One replicate-block reportable result was generated at sequence start and one at sequence end, each defined as the mean of three technical injections. Acceptance limits were derived from L1 validation performance. SIL-HCP recovery for each bracket was required to fall within 70–130% of nominal. Failure of the opening bracket precluded sequence initiation; failure of the closing bracket following a passing opening bracket triggered deviation investigation prior to result release.

An orthogonal SST component employed a retention time calibration (RTC) peptide mixture injected at sequence start and end. Two metrics were monitored: median retention time deviation (ΔRT) relative to a locked deployment reference and median RTC MS1 signal intensity relative to the Phase I baseline median. Acceptance thresholds of ±0.50 min (ΔRT) and ±35% (MS1 intensity) were implemented during initial operational deployment and remain subject to refinement based on Phase I control-chart performance.

Longitudinal monitoring of run-level SST metrics was performed using Phase I Individuals–Moving Range (I–MR) control charts. Control-limit derivation and monitoring procedures are described in **Supplementary Method S3**. Quantitative SIL-HCP limits are directly traceable to the validated TE performance envelope.

## 3. Results

Validation outcomes for specificity, linearity, trueness, precision, range, and limits of quantifications are presented under the predefined TE decision criteria.

### 3.1 Protein inference and identification depth

Protein inference was performed using a deterministic greedy parsimony algorithm applied to the theoretical tryptic digest of the reference FASTA. Vendor-reported and parsimony-derived group sizes showed substantial concordance along the identity line (**Figure 2A**). Discordance was concentrated among vendor-defined singleton and small groups (<10 proteins), which expanded into modest multi-protein equivalence classes under parsimony rules. Agreement increased with group size, reflecting rule-dependent resolution of shared-peptide networks.

**Figure 2.**
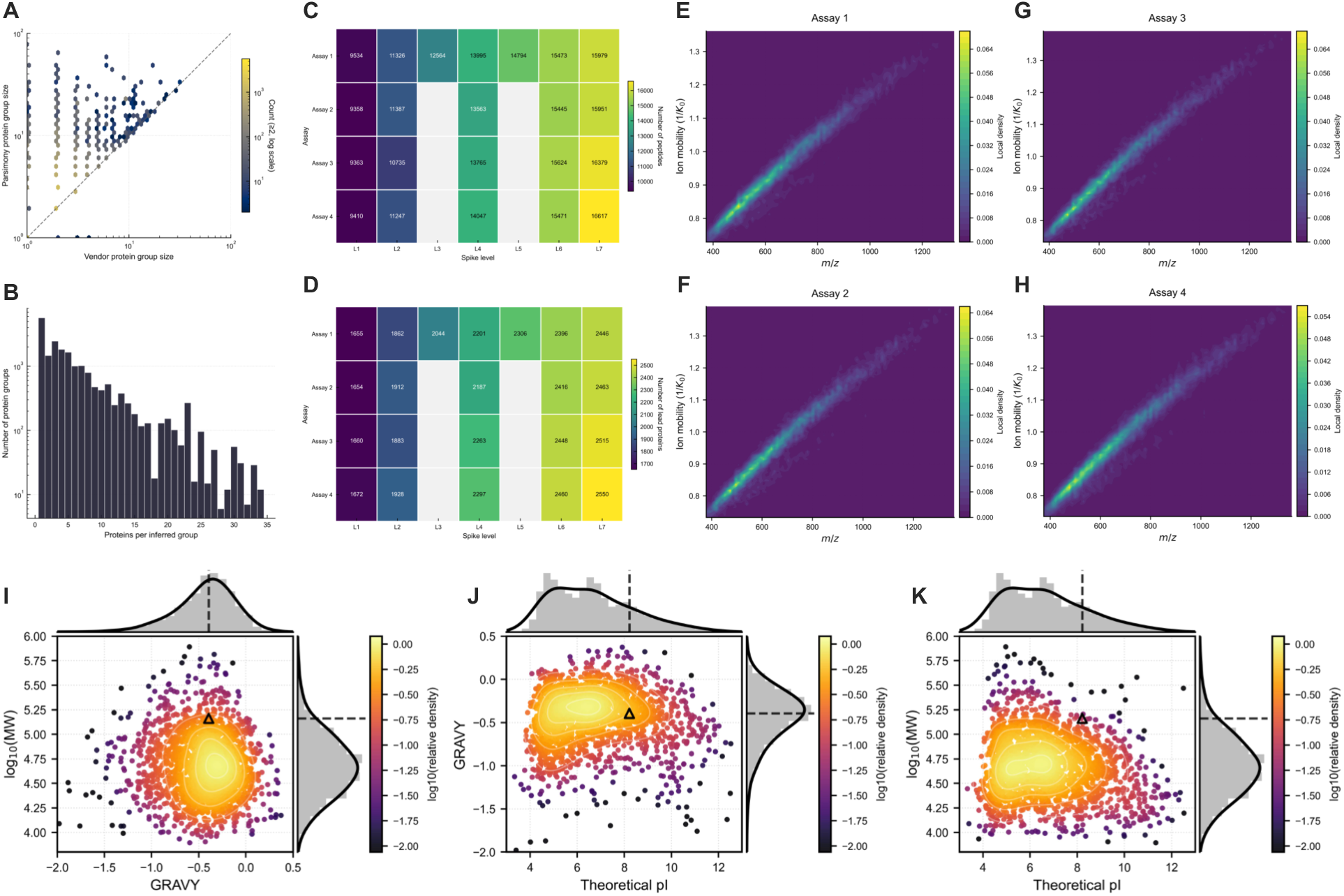
(**A**) Concordance between vendor-reported and parsimony-derived protein group sizes (Hexbin, log–log scale; dashed line = identity). (**B**) Distribution of parsimony-derived group sizes. (**C–D**) Peptide and lead-protein counts across assays and spike levels. (**E–H**) Precursor occupancy in the m/z–1/K₀ plane for each assay, shown as density-weighted scatter plots. (**I–K**) Physicochemical distribution of inferred CHO proteins (GRAVY, pI, log₁₀MW); the NISTmAb reference is highlighted.

Parsimony inference yielded a broad distribution of group sizes (**Figure 2B**), extending to approximately 25–30 proteins per group. Group frequency decreased monotonically with increasing size. Singleton groups predominated, but multi-member equivalence classes constituted a structured fraction of the inferred proteome, consistent with non-unique peptide connectivity under theoretical digestion.

In Assay 1, unique peptides increased from 9,534 (L1) to 15,979 (L7), and inferred lead proteins from 1,655 to 2,446 (**Figure 2C–D**). Assays 2–4 produced closely matched counts at shared levels. At 50 ng (L4), peptide identifications ranged from 13,563–14,047 and lead proteins from 2,187–2,297 (inter-assay CV <2% and <3%, respectively). At 20 ng, counts spanned 9,358–9,410 peptides and 1,654–1,672 proteins.

Precursor-space reproducibility was examined in the m/z–1/K₀ plane (**Figure 2E–H**). All assays exhibited the characteristic diagonal density distribution defined by mass–collisional cross-section coupling. Kernel density maxima overlapped without measurable displacement, confirming invariant precursor occupancy across runs. Variation in identification depth was attributable to analyte load, not acquisition instability.

Physicochemical coverage of inferred CHO proteins was characterized by theoretical pI, GRAVY index, and log₁₀(MW) (**Figure 2I–K**). pI values ranged from <4 to >12 with maximal density between 5–7. GRAVY values spanned −2.0 to +0.5 (centered ∼−0.6). MWs extended from <10 kDa to >500 kDa with modal density between 30–100 kDa. The NIST mAb occupied a peripheral

Spearman protein–protein correlation matrices were computed from log₁₀-transformed intensities. Average-linkage clustering of (1 − ρ) dissimilarities was applied to the 1,250 proteins with the highest inter-sample variance at each spike level (**Figure 3A**). Silhouette maximization over k = 2–12 selected k = 2 at all levels; silhouette widths ranged from 0.237 to 0.285, below conventional thresholds for meaningful cluster separation. Cophenetic correlation coefficients ranged from 0.526 to 0.600 (**Supplementary Table S1, Panel A**) with no systematic dependence on HCP load. The binary partition lacks compositional coherence and does not define a clearly interpretable grouping.

**Figure 3.**
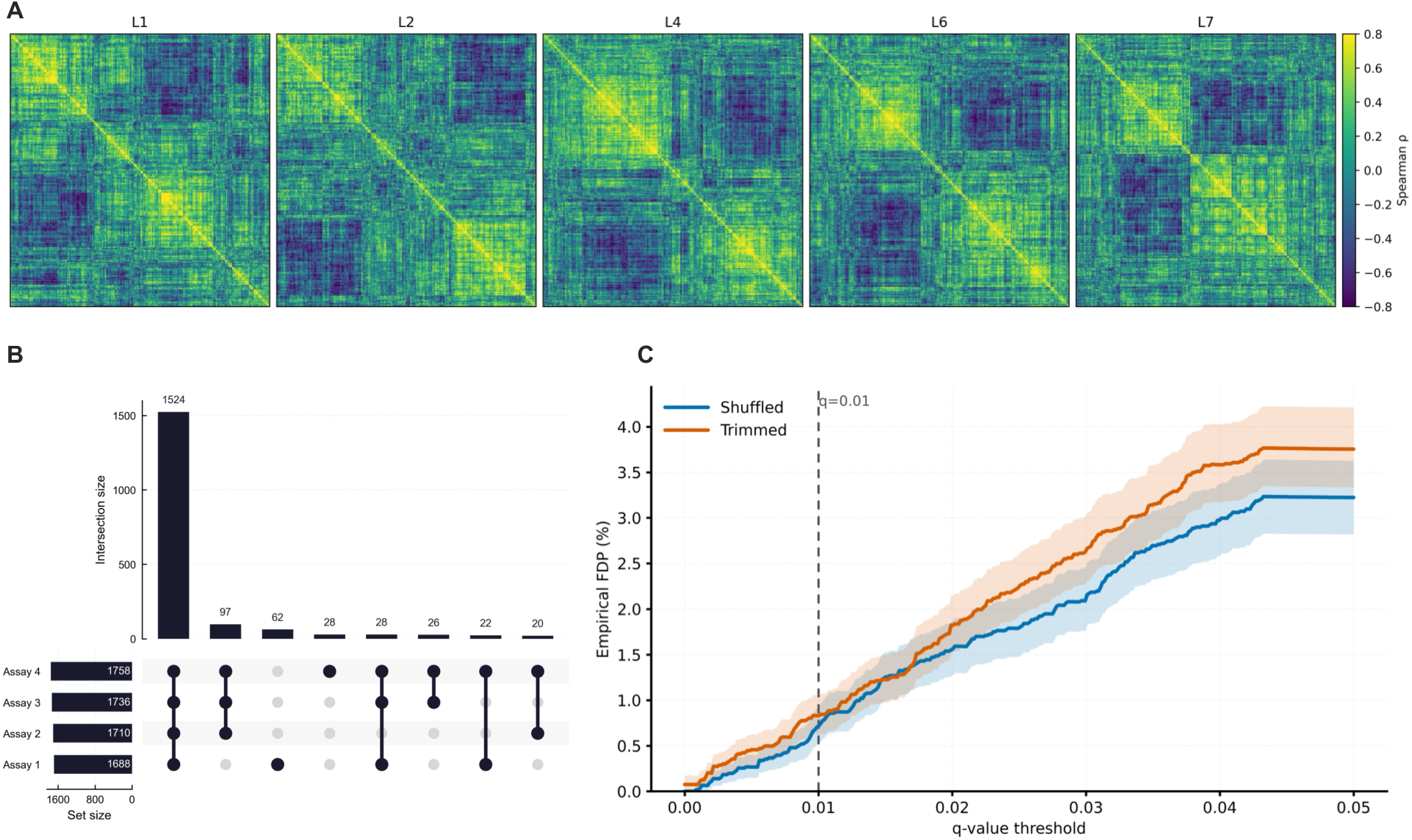
(**A**) Spearman protein–protein correlation matrices over common spike levels (L1, L2, L4, L6, L7), hierarchically clustered on a fixed correlation scale. (**B**) UpSet plot of protein identification overlap across the four assays. (**C**) Empirical peptide-level false discovery proportion (FDP) as a function of q-value threshold (τ) estimated by search-embedded entrapment; shaded regions denote 95% bootstrap bands and the dashed line marks q = 0.01.

Matrix similarity was evaluated on 648 proteins common to all spike levels using Mantel permutation tests (Supplementary Table S1, Panel B). All ten pairwise comparisons rejected matrix independence (p < 0.001). Mantel r values were small (0.062–0.298) and decreased with increasing spike-level separation. The protein–protein correlation structure lacks stability across concentration levels and does not justify structured covariance modeling.

The absence of discrete covariance structure supports the exchangeability assumption underlying the one-way random-effects restricted maximum likelihood (REML) variance decomposition. More complex covariance specifications, including cluster-indexed random effects or block-diagonal residual covariance models, were not required for this dataset.

Consistency of quantifiable proteins was assessed by UpSet analysis (**Figure 3B**). A total of 1,524 proteins were quantified in all four assays (87–90% of assay-specific inventories: 1,688–1,758 proteins). Three-assay intersections comprised 97 and 62 proteins; assay-unique identifications were minimal.

### 3.2 Specificity and False Discovery Control

No SIL-HCP heavy peptides were detected in the unspiked mAb control (L0). Endogenous light CHO peptides do not constitute a blank failure mode because quantification is restricted to the isotopically resolved SIL-HCP population.

Peptide-level specificity was evaluated using search-embedded entrapment with concatenated target and entrapment databases constructed to approximate a 1:1 ratio. Empirical FDP curves with pointwise 95% bootstrap percentile bands are shown in **Figure 3C**. FDP increased as the reported q-value threshold was relaxed. At the operating threshold (q = 0.01), empirical FDP was below 1% for both entrapment constructions: 0.7–0.8% (shuffled) and 0.85–0.9% (trimmed).

The trimmed entrapment space produced marginally higher FDP at stringent thresholds, consistent with increased score competitiveness from preserved amino acid composition. At q = 0.01, both constructions converged to comparable empirical control. Bootstrap-derived FDP provides the primary calibration of identification error. The two-sided 95% Wilson score interval for the entrapment proportion at q = 0.01 (**Supplementary Table S2**) was consistent with binomial sampling variability.

Hi3 quantification requires three concordant peptides per protein group. Assuming approximate independence and peptide-level FDP ≈ 0.009, the probability of three false-positive peptides mapping to the same protein group is on the order of 10⁻⁷. Protein-level false identification is therefore negligible relative to other variance components in the validation; a formal derivation is provided in **Supplementary Note S1**.

Blank-matrix evaluation excludes spike carryover and matrix-driven false positives. Entrapment analysis provides an empirical upper bound on peptide-level identification error independent of target–decoy competition. Specificity at the reporting threshold is therefore supported.

### 3.3 Linearity

WLS regression yielded the fitted relationship y = 1.25 + 0.798x, R^2^ = 0.9928 (**Figure 4A**; **Table 2**). The slope was below unity (HC3 95% CI [0.79, 0.81]; p < 0.001), corresponding to proportional compression of approximately 20% over 20–80 ng. The intercept was positive (HC3 95% CI [0.83, 1.67] ng; p < 0.001), introducing an additive offset whose relative contribution decreased with concentration (∼7% at 20 ng; <2% at 80 ng).

**Figure 4.**
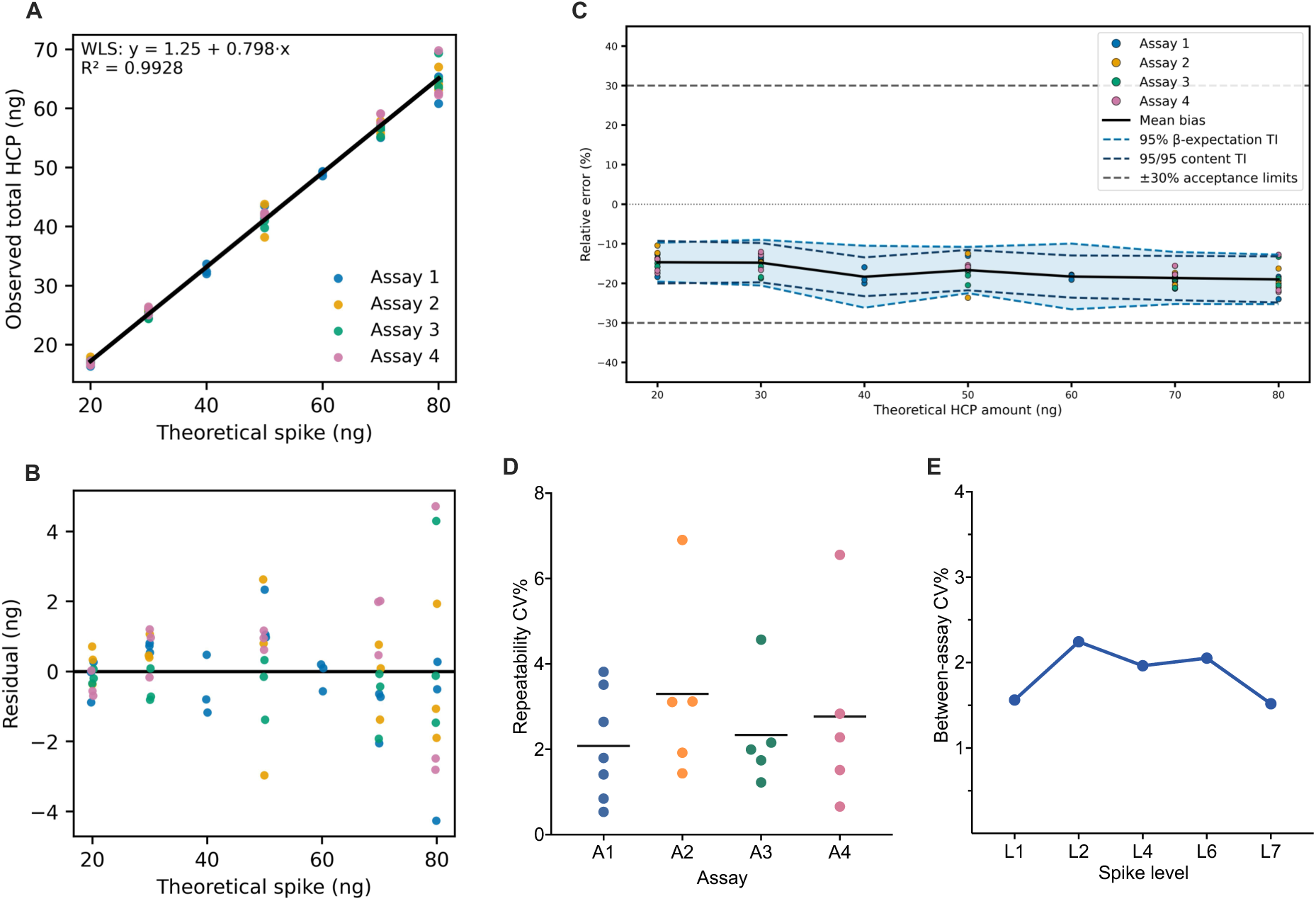
(**A**) WLS regression of observed versus nominal HCP amounts (20–80 ng). (**B**) Regression residuals across spike levels. (**C**) Accuracy profile on the relative-error scale showing mean bias (solid line), 95% β-expectation tolerance limits (blue dashed), and 95/95 content tolerance limits (green dashed); horizontal lines denote ±30% acceptance criteria. (**D**) Within-assay repeatability (CV%) by spike level. (**E**) Between-assay dispersion (CV%) by spike level.

**Table 2.** Linearity summary for total HCP quantification.

| Metric | Value |
| --- | --- |
| Intercept (ng) | 1.25 |
| Intercept SE (HC3) | 0.21 |
| Intercept 95% CI (HC3) | [0.83-1.67] |
| Slope | 0.7978 |
| Slope SE (HC3) | 0.01 |
| Slope 95% CI (HC3) | [0.79-0.81] |
| R<sup>2</sup> | 0.9928 |
| P (Intercept = 0) | < 0.001 |
| P (Slope = 1) | < 0.001 |

The implied relative bias, 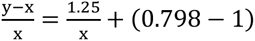, approaches −20.2% as x → ∞, with partial attenuation at lower concentrations due to the intercept term.

Residuals were centered without curvature over the tested range (**Figure 4B**). Absolute dispersion increased with concentration, consistent with intensity-dependent variance in label-free quantification. Inverse-variance weighting stabilized residual variance. Replicate-block responses from all assays followed a common regression function without assay-specific displacement. Compression and offset therefore represent stable properties of the measurement system and were incorporated into subsequent trueness and TE analyses.

### 3.4 Trueness

Trueness was evaluated at the replicate-block level as the signed deviation of mean total HCP from the nominal spike amount (**Table 3**). Bias was negative across 20–80 ng, ranging from −14.67% at 20 ng to −19.01% at 80 ng, corresponding to mean recoveries of 85.33% to 80.99%. Bias magnitude increased between 20 and 40 ng and stabilized between −16.7% and −19.0% from 50 to 80 ng. The P05–P95 distribution of replicate-block relative errors showed consistent negative centering at all levels. The 95% confidence interval (CI) for recovery remained below 100% throughout the range.

**Table 3.**
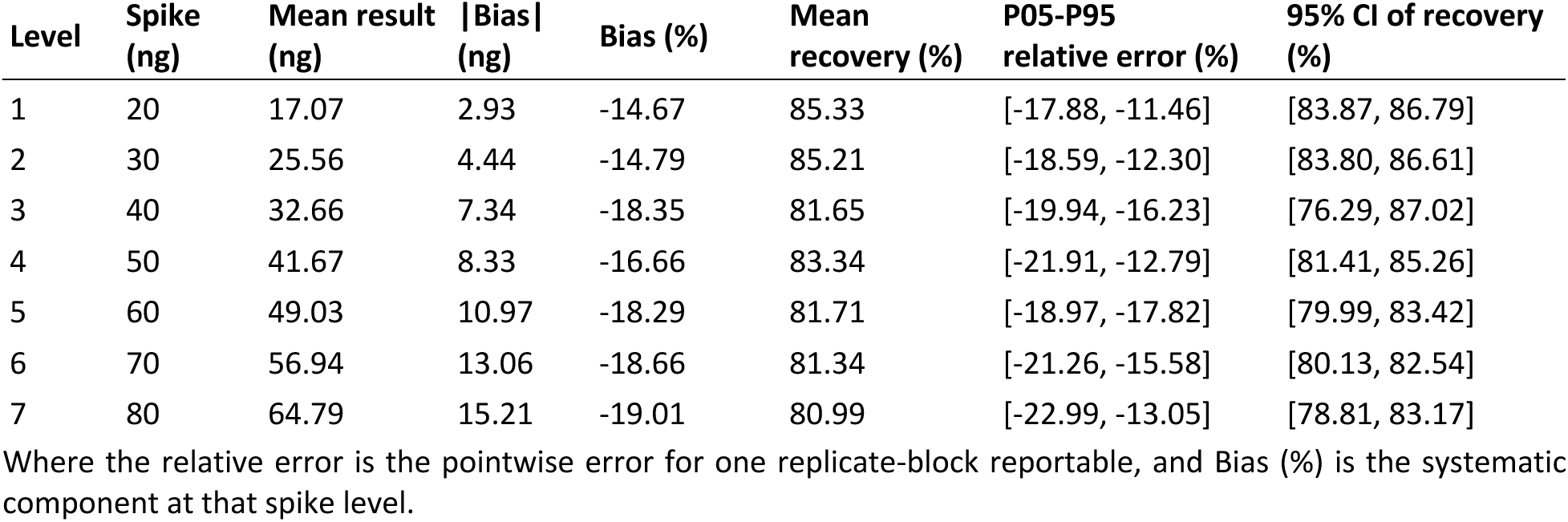
Trueness summary for total HCP quantification.

The bias trajectory aligns with the fitted calibration function. Relative bias becomes increasingly negative with concentration and approaches the asymptotic compression implied by the slope deficit. At 80 ng, the model-implied bias (−18.6%) closely matched the empirical estimate (−19.01%). The trueness profile therefore reflects stable proportional compression and was incorporated into total-error evaluation.

### 3.5 Precision

Precision was evaluated on the replicate-block relative-error scale using a one-way random-effects ANOVA with assay as the grouping factor (**Table 4**). Variance components were estimated using method-of-moments estimators. Repeatability (within-assay SD, 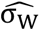) dominated at all spike levels and ranged from 0.69% (60 ng) to 3.81% (80 ng), with most values between 1% and 3%.

**Table 4.** Precision summary for total HCP quantification.

| Level | Spike (ng) | Mean result (ng) | Within-assay SD (%) | Between-assay SD (%) | Total SD (%) | Within-assay df | Between-assay df |
| --- | --- | --- | --- | --- | --- | --- | --- |
| 1 | 20 | 17.07 | 2.29 | 0.14 | 2.30 | 8 | 3 |
| 2 | 30 | 25.56 | 1.61 | 1.67 | 2.32 | 8 | 3 |
| 3 | 40 | 32.66 | 2.16 | 0 | 2.16 | 2 | 0 |
| 4 | 50 | 41.67 | 3.09 | 0 | 3.09 | 8 | 3 |
| 5 | 60 | 49.03 | 0.69 | 0 | 0.69 | 2 | 0 |
| 6 | 70 | 56.94 | 1.35 | 1.48 | 2.00 | 8 | 3 |
| 7 | 80 | 64.79 | 3.81 | 0 | 3.81 | 8 | 3 |

Variance components were estimated by one-way random-effects ANOVA using method-of-moments estimators. For levels present in a single assay (40 and 60 ng), between-assay variance was not estimable. Total SD corresponds to intermediate precision under ICH Q2(R2).

The between-assay component 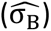 was small and was truncated to zero when MS_between_ ≤ MS_within_. Non-zero estimates were observed at 30 ng (1.67%) and 70 ng (1.48%), without systematic concentration dependence.

Total SD 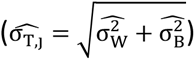, corresponding to intermediate precision, ranged from 0.69% to 3.81%, with the largest values at 50 ng (3.09%) and 80 ng (3.81%) (Table 4). Over the validated range, total variability was primarily driven by within-assay dispersion.

Median repeatability CVs across assays were from 2.1% to 3.2% (**Figure 4D**). Between-assay CV ranged from 1.5% to 2.3% (**Figure 4E**).

Levels 40 ng and 60 ng were present in Assay 1 only; between-assay variance was therefore non-estimable (df = 0). Precision estimates at these levels reflect repeatability only and contribute to the variance model through the within-assay component.

For levels present in all assays (between-assay df = 3), the hierarchical design limits detection of small between-assay components. The minimum detectable between-assay SD at 80% power is approximately 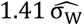 (**Supplementary Note S2**; **Table S3**). At median 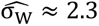, between-assay SD below ∼3.2% is statistically indistinguishable from zero. Method-of-moments truncation to zero should therefore be interpreted as compatibility with small between-assay variance and not absence of such variance. REML estimates (**Supplementary Table S4**) yielded small positive components where truncation occurred, supporting robustness of the variance decomposition.

### 3.6 Total-error accuracy profile

Accuracy was evaluated on the relative-error scale within the predefined TE approach. Across 20–80 ng, bias was negative at all levels (−14.67% to −19.01%) and exceeded the random component in magnitude. Model-predicted total SD ranged from 2.22% to 2.72%, indicating dominance of systematic under-recovery.

Level-specific 95% β-expectation TIs, incorporating Satterthwaite-adjusted effective degrees of freedom (3.0–10.78), ranged from [−19.57, −9.76]% at 20 ng to [−25.28, −12.74]% at 80 ng (**Table 5**; **Figure 4C**). All β-expectation TIs lay entirely below 0%. Interval width increased modestly with concentration, consistent with the log–log variance model.

**Table 5.** Accuracy summary for total HCP quantification.

| Level | Spike (ng) | Mean result (ng) | Bias (%) | Pred. total SD (%) | Satterthwaite df | 95 β-expectation TI (%) | 95/95 content TI (%) |
| --- | --- | --- | --- | --- | --- | --- | --- |
| 1 | 20 | 17.07 | –14.67 | 2.22 | 10.78 | [–19.57, –9.76] | [–20.03, –9.30] |
| 2 | 30 | 25.56 | –14.79 | 2.36 | 6.02 | [–20.56, –9.03] | [–19.76, –9.83] |
| 3 | 40 | 32.66 | –18.35 | 2.46 | 3 | [–26.17, –10.52] | [–23.26, –13.43] |
| 4 | 50 | 41.67 | –16.66 | 2.54 | 8 | [–22.52, –10.81] | [–21.77, –11.56] |
| 5 | 60 | 49.03 | –18.29 | 2.61 | 3 | [–26.59, –9.99] | [–23.63, –12.96] |
| 6 | 70 | 56.94 | –18.66 | 2.67 | 5.77 | [–25.25, –12.07] | [–24.25, –13.08] |
| 7 | 80 | 64.79 | –19.01 | 2.72 | 8 | [–25.28, –12.74] | [–24.84, –13.18] |

Predicted total SD denotes the smoothed estimate on the relative-error scale. Effective degrees of freedom were obtained via Welch–Satterthwaite adjustment.

Parallel 95/95 content TIs obtained by hierarchical cluster bootstrap were comparable in location and width, spanning [−20.03, −9.30]% at 20 ng to [−24.84, −13.18]% at 80 ng.

Both TI types were fully contained within the predefined ±30% acceptance limits at all spike levels. The most conservative case occurred at 80 ng, where the lower bound of the 95/95 content TI (−24.84%) remained 5.16 percentage points above the −30% criterion.

### 3.7 Analytical Range, LOD, and LLOQ

The analytical range was defined using the TE criterion requiring the 95% β-expectation TI for replicate-block relative error to lie within ±30%. This condition was satisfied at all tested levels from 20 to 80 ng (**Table 5**; **Figure 4C**). At 80 ng, the lower β-expectation bound (−25.28%) remained 4.72 percentage points within the −30% acceptance limit. The validated analytical range is therefore 20–80 ng total HCP per injection.

Consistent with ICH Q2(R2) Section 6, the LLOQ was defined as the lowest experimentally tested level meeting the β-expectation criterion. Accordingly, 20 ng constitutes the validated LLOQ. The corresponding 95/95 content TI provided confirmatory coverage. Levels below 20 ng were not evaluated, and no extrapolation beyond the validated domain was performed.

A signal-to-noise–based LOD is not applicable to this estimator. Total HCP is derived from aggregated protein-group signals under a calibration-compression model; detectability is governed by population-level quantitative performance rather than single-analyte intensity. Reporting is restricted to the validated TE range.

### 3.8 Abundance-stratified total-error performance

Total HCP is an aggregate estimator over a heterogeneous abundance distribution. Aggregation reduces variance through averaging but can obscure abundance-dependent bias. TE performance was therefore re-estimated within fixed abundance strata (Q1–Q4) defined at L4. Bootstrap KDEs were well separated by spike level within each stratum (**Figure 5A**); with 10,000 resamples per cell, bootstrap variability was negligible relative to between-level separation.

**Figure 5.**
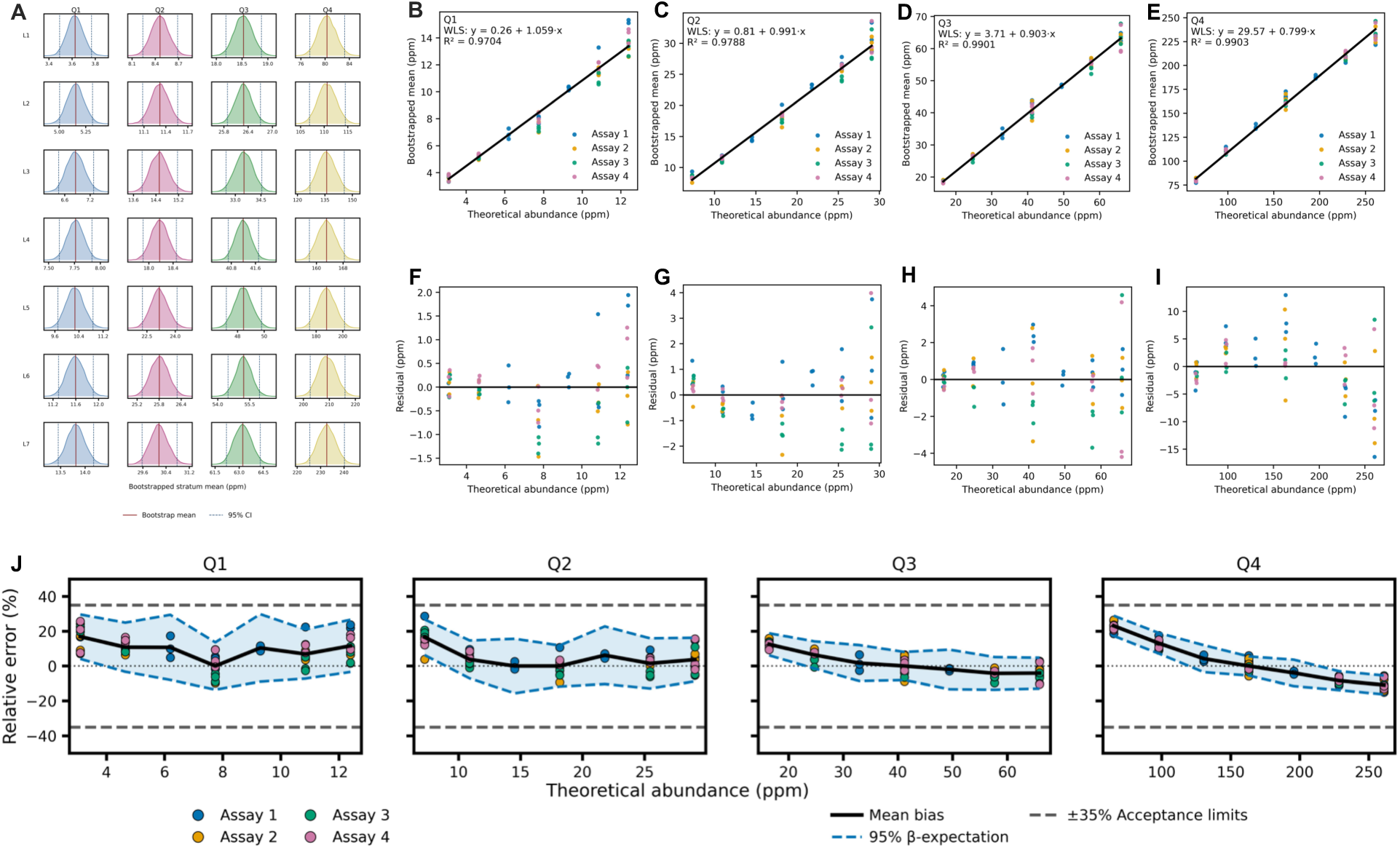
(**A**) (A) Bootstrap density distributions of stratum-level mean abundances (ppm; 10,000 resamples) across spike levels for Q1–Q4; solid lines denote bootstrap means and dashed lines the 2.5th–97.5th percentiles. (**B–I**) Weighted least-squares calibration of bootstrapped mean abundance versus theoretical abundance for strata Q1–Q4; upper panels show fitted relationships and lower panels residuals. (**J**) Stratified accuracy profiles showing relative error of observed-to-anchor response ratios versus theoretical spike ratios; solid lines indicate mean bias, and dashed lines denote 95% β-expectation TIs with ±35% acceptance criteria.

Stratified WLS regressions (**Table 6**; **Figure 5B–E**) demonstrated abundance-dependent calibration behavior. Slopes decreased with abundance rank: Q1 = 1.06 (95% CI [1.02, 1.10]), Q2 = 0.99 [0.95, 1.04], Q3 = 0.90 [0.89, 0.92], Q4 = 0.80 [0.78, 0.81]. Low-abundance proteins exhibited mild expansion, mid-abundance proteins approximated proportionality, and higher-abundance strata exhibited compression. Intercepts increased with abundance rank (0.26 → 29.57 ppm), introducing positive offsets at low expected abundance. Residual diagnostics showed no curvature (**Figure 5F–I**).

**Table 6.** Linearity summary for abundance-stratified HCP quantification.

| Metric | Value Q1 | Value Q2 | Value Q3 | Value Q4 |
| --- | --- | --- | --- | --- |
| Intercept (ppm) | 0.26 | 0.81 | 3.71 | 29.57 |
| Intercept SE (HC3) | 0.09 | 0.27 | 0.18 | 0.92 |
| Intercept 95% CI (HC3) | [0.07-0.44] | [0.27-1.36] | [3.35-4.07] | [27.73-31.41] |
| Slope | 1.0588 | 0.9906 | 0.9029 | 0.7986 |
| Slope SE (HC3) | 0.02 | 0.02 | 0.01 | 0.01 |
| Slope 95% CI (HC3) | [1.02-1.10] | [0.95-1.04] | [0.89-0.92] | [0.78-0.81] |
| R2 | 0.9704 | 0.9788 | 0.9901 | 0.9903 |
| p(Intercept = 0) | <0.01 | <0.001 | <0.001 | <0.001 |
| p(Slope = 1) | <0.001 | <0.001 | <0.001 | <0.001 |

Trueness patterns followed from these calibration structures (**Supplementary Table S6**). Q1 and Q2 displayed positive bias at most levels, whereas Q3 and Q4 transitioned from positive bias at low spike ratios to negative bias at higher ratios. In Q4, bias ranged from +23.1% (L1) to −10.9% (L7), consistent with slope compression below unity combined with positive intercept offset.

Stratified bias is defined relative to the L4 anchor. Positive bias in Q1 reflects proportional deviation from the L4 systematic component and does not imply absolute over-recovery relative to nominal concentration. Absolute bias equals the aggregate L4 bias plus the stratum-specific deviation (**Supplementary Note S3**).

Variance decomposition (**Supplementary Table S7**) attributed most dispersion to repeatability. Within-assay SDs were largest in Q1 (up to 8.06%) and decreased with abundance rank. Between-assay components were smaller and frequently truncated to zero under the non-negativity constraint. Total SD ranged from 0.80% (Q3, L5) to 8.06% (Q1, L1). Elevated dispersion in Q1 reflects higher leverage of individual proteins within sparse strata.

Stratum-specific TE accuracy profiles (**Supplementary Table S8**; **Figure 5J**) were fully contained within ±35% acceptance limits at all spike levels. Interval displacement was driven primarily by bias. The widest β-expectation interval occurred in Q1 at L1 [6.0, 27.7]%, and the most negative in Q4 at L7 [−15.5, −6.3]%.

The abundance-aware validated range was defined as the intersection of acceptable spike levels across Q1–Q4. The lowest passing level in Q1 defined the abundance-aware LLOQ (mean 3.62 ppm; conservative P95 = 3.87 ppm). The highest passing level in Q4 defined the abundance-aware ULOQ (mean 232.57 ppm; conservative P05 = 222.97 ppm).

The aggregate LLOQ (20 ng per injection) differs from the abundance-aware LLOQ (3.6 ppm; conservative P95 = 3.87 ppm). The former defines the minimum validated total impurity burden at the sample level, whereas the latter defines the lowest abundance class with demonstrated population-level quantitative performance. These limits address different measurement constructs and are not interchangeable (**Supplementary Table S5**).

REML estimates (**Supplementary Table S9**; **Figure S1**) yielded small positive between-assay components where method-of-moments truncation occurred, supporting robustness of the variance decomposition.

The aggregate total HCP remains the validated reportable quantity for impurity burden assessment. Abundance-stratified analysis characterizes performance heterogeneity without redefining the primary release measurand.

### 3.9 Robustness

Robustness was evaluated by varying two analytical factors: data-processing workflow and MS platform. Total HCP (ng) was the reportable. Agreement was assessed using Deming regression (2,000 bootstrap resamples for 95% CIs), Lin’s concordance correlation coefficient (CCC), and Bland–Altman analysis on the relative-difference scale. Pairwise 95% limits of agreement (LoA) were evaluated against the predefined ±30% acceptance limits.

#### 3.9.1 Software comparison: SpectroMine vs FragPipe

A subset of Assay 1 timsTOF Pro data (L1–L7; one preparation per level; technical triplicates retained) was reprocessed using FragPipe (v24.0; MSFragger v4.4.1; IonQuant v1.11.20). Database composition, FDR thresholds, and Hi3 quantification parameters were held constant. Deming regression of FragPipe versus SpectroMine yielded slope 0.959 (95% CI [0.943, 0.984]) and intercept +1.80 ng (**Figure 6A**; **Table 7**). Lin’s CCC was 0.9982. Bland–Altman analysis showed mean relative bias +1.27% with 95% LoA [−5.8, +8.3]% (**Figure 6B**). All individual differences and corresponding LoA were within ±30%.

**Figure 6.**
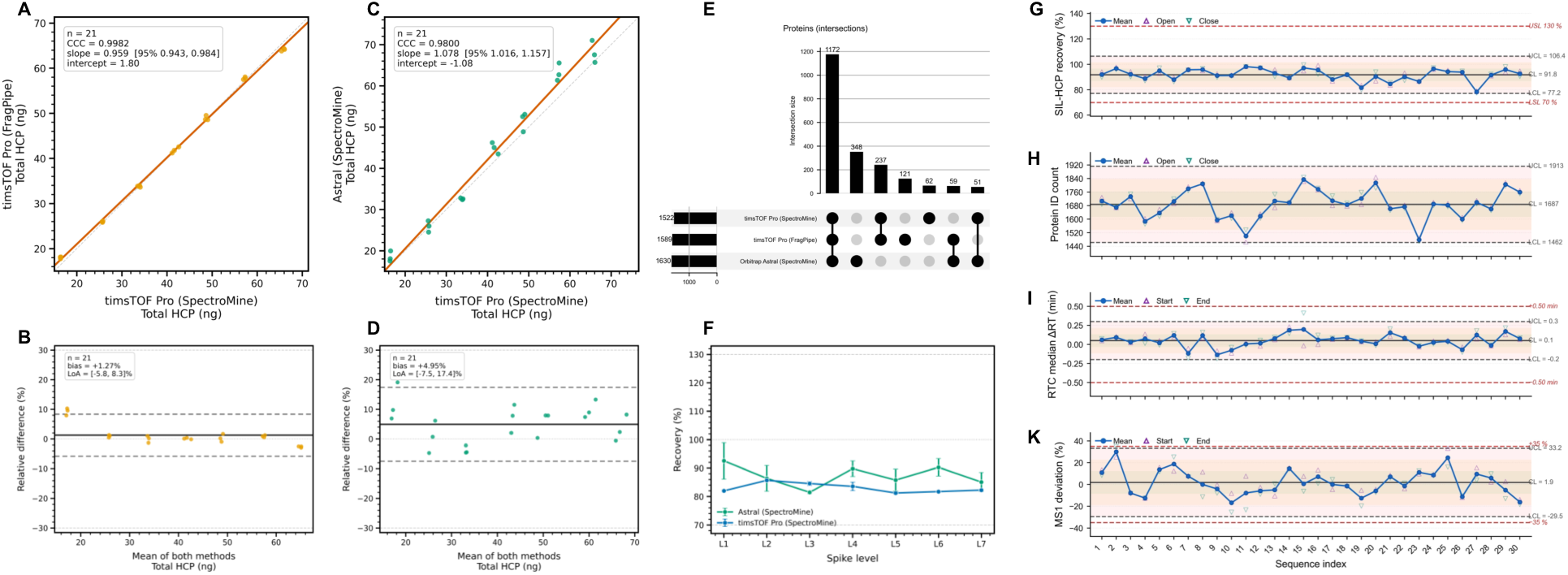
(**A**) Deming regression of total HCP quantified by FragPipe versus SpectroMine on identical timsTOF Pro data; dashed line indicates identity. (**B**) Bland–Altman analysis for software comparison. (**C**) Deming regression of total HCP between timsTOF Pro and Orbitrap Astral (SpectroMine processing). (**D**) Bland–Altman analysis for cross-platform comparison. (**E**) UpSet plot showing overlap of quantified proteins across analytical configurations. (**F**) Level-wise mean recovery (%) for timsTOF Pro and Orbitrap Astral; dashed lines denote 70% and 130% acceptance limits. (**G–J**) Phase I Individuals control charts for run-level SST metrics: SIL-HCP recovery, protein identification count, RTC median retention time deviation, and RTC MS1 intensity deviation; red dashed lines indicate specification limits. (**K**) Corresponding moving-range charts showing short-term variability.

**Table 7.** Agreement statistics for robustness comparisons.

| Category | Comparison | Deming slope<br>[95% CI] | Deming<br>intercept<br>(ng) | Lin's CCC | BA bias<br>(%) | BA LoA<br>(%) | Within<br>±30% |
| --- | --- | --- | --- | --- | --- | --- | --- |
| Software | SpectroMine vs<br>FragPipe | 0.9592 [0.9433, 0.9843] | +1.805 | 0.9982 | 1.27 | [-5.8, +8.33] | Yes |
| Platform | timsTOF Pro vs<br>Astral | 1.0785 [1.0158, 1.1569] | -1.081 | 0.9800 | 4.95 | [-7.51, +17.4] | Yes |

#### 3.9.2 Platform comparison: timsTOF Pro vs Orbitrap Astral

Total HCP values from timsTOF Pro and Orbitrap Astral were compared following identical SpectroMine processing and matched 500 ng injection load (L1–L7; 20–80 ng). Deming regression yielded slope 1.078 (95% CI [1.016, 1.157]) and intercept −1.08 ng (**Figure 6C**; **Table 7**). Lin’s CCC was 0.9800. Bland–Altman analysis showed mean relative bias +4.95% with 95% LoA [−7.5, +17.4]% (**Figure 6D**). The upper LoA bound remained below the +30% acceptance limit. All individual differences were contained within ±30%.

Quantifiable protein overlap across SpectroMine (timsTOF Pro), FragPipe (timsTOF Pro), and SpectroMine (Orbitrap Astral) is shown in **Figure 6E**. A core set of 1,172 proteins was shared across configurations, with smaller configuration-specific subsets.

Level-wise mean recovery (**Figure 6F**) ranged from 81–85% on timsTOF Pro and 85–99% on Orbitrap Astral. LoAs were narrower for software reprocessing than for platform comparison. Observed pairwise differences were of similar magnitude to the estimated intermediate precision from hierarchical validation.

Under ICH Q14 lifecycle principles, platform transfer requires bridging validation demonstrating preservation of variance structure, not solely aggregate agreement.

### 3.10 System suitability Testing and process stability

SST performance was evaluated across 30 pilot analytical sequences using Phase I Individuals–Moving Range (I–MR) charts (**Figure 6G–K**; **Supplementary Figure S2**). Run-level SIL-HCP recovery remained within the 70–130% specification limits for all sequences and was centered near the baseline mean without sustained 3σ violations. Moving-range charts showed no persistent inflation of short-term variability.

Protein identification counts exhibited expected stochastic dispersion associated with DDA. Isolated downward excursions were observed but were not sustained and did not coincide with recovery failure or chromatographic deviation. Moving-range values showed intermittent spikes without evidence of process drift.

RTC median retention time deviation remained within the ±0.50 min specification limits and centered on the deployment reference. RTC MS1 deviation remained within the ±35% tolerance. A single elevated MS1 observation was not accompanied by concurrent recovery or ΔRT shifts and was not followed by sustained deviation. Moving-range charts indicated stable short-term variability for recovery and ΔRT, with intermittent variability in identification count and MS1 deviation consistent with operational noise.

Across all monitored metrics, no sustained loss of statistical control or specification-limit exceedance was observed.

## 4. Discussion

This validation demonstrates that label-free ddaPASEF proteomics, evaluated within the TE approach, satisfies prespecified performance criteria aligned with ICH Q2(R2) for quantitative HCP determination within 20–80 ng. To our knowledge, this is the first prospective validation of an untargeted proteomic workflow for HCP quantification conducted under ICH Q2(R2) guidelines. The approach validates the measurement system itself, encompassing identification, inference, and quantitative aggregation steps that jointly determine the reportable measurand, in contrast to targeted MS assays that define analytes a priori.

Systematic and random error components were modeled separately and incorporated into predictive TI constructions. All tested levels satisfied containment under both 95% β-expectation and 95/95 content TI definitions. The WLS calibration model (ŷ = 1.25 + 0.798x, R^2^ = 0.993) comprises a fixed additive offset contributing a bias term that decays as 1/x, and a proportional compression term (slope deficit ≈0.20) inducing concentration-independent multiplicative attenuation. HC3-robust inference confirmed deviation from β₀ = 0 and β₁ = 1; residual diagnostics supported first-order specification without curvature. The response structure was reproducible within and between four independent assays. The ±30% aggregate acceptance limit was derived from the USP ⟨1132⟩ immunoassay performance envelope and SFSTP β-expectation guidance.

Precision was low relative to bias magnitude. Median repeatability was 2.7% CV; intermediate precision ranged from 1.5% to 2.3% CV. Between-assay variance was minor and frequently truncated to zero under the method-of-moments estimator. Two features of the measurement system account for this: the locked acquisition method yielded stable precursor-space occupancy and reproducible identification depth, and total HCP, as a composite estimator, attenuates independent protein-level fluctuations upon summation in proportion to population size. With df = 3 for the between-assay component, between-assay SD below approximately 3.2% is statistically indistinguishable from zero (**Supplementary Note S2**). Zero estimates therefore reflect limited resolving power at the available degrees of freedom, not absence of between-assay variability. Log–log variance modeling confirmed near-constant relative-error dispersion within the validated domain.

The method exhibits a stable negative bias of approximately −15% to −19%, with mean recoveries between 80.99% and 85.33%. This under-recovery corresponds to the proportional compression in the calibration function and arises from three multiplicative effects: incomplete proteolytic conversion for structurally resistant proteins at the fixed enzyme-to-substrate ratio; ionization suppression within the mAb-dominated matrix; and response-factor mismatch between the MassPREP Hi3 standards and CHO peptides, compounded by residual response-factor dispersion in the SIL-HCP calibrant proteome (**Supplementary Table S11**). The missed-cleavage distribution (95.81% zero, 4.19% one; **Supplementary Table S10**) falls within standard bottom-up benchmarks, implying that digestion efficiency is not the dominant contributor. Conservative peptide filtering introduces an additional bias–precision trade-off: exclusion of single-hit, high-variance, and incompletely detected peptides restricts Hi3 quantification to reproducible precursors per protein group. Each reportable result corresponds to the arithmetic mean of three injections, retaining only peptides detected in all acquisitions with acceptable CV. This suppresses stochastic DDA sampling noise at the peptide level but decreases the number of contributing proteins, attenuating recovered total HCP abundance. The attenuation is absorbed into the calibration model as part of the slope deficit and is evaluated jointly with all other systematic components within the TE construction. The ∼20% aggregate compression measured here falls within published Hi3 accuracy benchmarks in complex biological matrices, where individual protein-level deviations of 30–50% are common^27,45^.

Between-assay CV of bias was <3% and systematic bias dominated the TE budget, exceeding total SD by 5–7-fold. Under ICH Q2(R2), zero bias is not required when systematic deviation is stable and incorporated into predictive limits. The 95% β-expectation and 95/95 content TIs incorporate the empirical bias at each level; at all validated levels the most conservative TI bound (−24.84% at 80 ng) remained more than 5 percentage points inside the −30% limit. The two constructions address different inferential targets: the former provides expected predictive coverage for a future replicate-block result; the latter provides confidence-guaranteed containment of at least 95% of future results. Concurrent containment under both criteria establishes predictive control under complementary definitions of coverage. Bias reduction would require correction factors derived from orthogonal quantification (**Supplementary Notes S3, S5**).

Peptide-level response-factor equivalence between the SIL-HCP calibrant and endogenous CHO HCPs has not been independently demonstrated, and the calibrant proteome inevitably differs in composition from the true endogenous impurity profile. Because the validated measurand is the aggregate total HCP mass derived from Hi3 summation over all quantifiable protein groups, individual protein-level response-factor deviations are partially averaged, and residual calibration mismatch is absorbed into the empirical bias term propagated through the TI construction. Published evaluations of MS-based HCP workflows confirm that heterogeneous protein mixtures can provide reliable aggregate HCP estimates despite imperfect proteome matching, provided systematic bias is empirically characterized within the validation framework^28,37^. This validation applies specifically to this SIL-HCP lot and processing configuration; a change in calibrant source or composition would require reassessment under the revalidation trigger matrix (**Supplementary Table S12**). A bridging study between alternative SIL proteome preparations remains a necessary extension.

Aggregate TE performance masks abundance-dependent heterogeneity. Stratified WLS regression produced slopes decreasing with abundance rank (Q1 ≈ 1.06; Q2 ≈ 0.99; Q3 ≈ 0.90; Q4 ≈ 0.80), with intercepts increasing correspondingly. The aggregate metric, dominated by Q4 species, reflects high-abundance protein behavior; low-abundance strata exhibit distinct calibration profiles obscured in the summed result. L4 normalization forces zero bias at the anchor level by construction; stratified bias represents deviation from L4-specific systematic error, not from ground truth. Absolute bias equals the aggregate L4 bias (−16.7%) plus the stratum-specific relative deviation (**Supplementary Note S3**). All strata satisfied ±35% β-expectation containment simultaneously, defining abundance-aware LLOQ and ULOQ expressed in ppm. The ±35% stratified acceptance limit accounts for additional variance from population-level bootstrap averaging within fixed abundance strata and stratum-to-stratum heterogeneity in calibration slope (**Supplementary Note S4**). The abundance-stratified LLOQ of 3.6 ppm (P95 = 3.87 ppm) provides additional resolution at the protein-population level. For individual high-risk protein tracking at sub-ppm concentrations, targeted LC–MS/MS methods with protein-specific internal standards remain the appropriate strategy.

Empirical entrapment analysis established peptide-level FDP below 1% at q = 0.01 under both shuffled and trimmed constructions, with bootstrap-derived FDP of 0.7–0.9% constituting the primary calibration evidence. Propagation to the protein level is attenuated geometrically by the Hi3 requirement of three concordant peptides: at FDP = 0.009, the expected protein-level false identification rate scales as FDP³ ≈ 10⁻⁷, negligible relative to other uncertainty components. Deterministic parsimony inference ensures denominator stability for longitudinal monitoring by preventing drift attributable to software-specific grouping heuristics. Cophenetic correlation analysis confirmed preservation of the protein–protein covariance structure at all spike levels, supporting the assumption that variance components are not confounded by concentration-dependent covariance shifts.

Perturbation of search engine (FragPipe) and MS platform (Orbitrap Astral) did not produce deviations exceeding ±30%. The wider platform limits of agreement reflect additional acquisition-layer variance from ion sampling architecture and duty cycle differences. Under ICH Q14, platform transfer constitutes a lifecycle change requiring bridging validation that demonstrates preservation of calibration behavior and variance structure, not solely aggregate agreement. The two-component SST framework addresses the gap between one-time validation and continued performance verification. LLOQ-proximal bracket injections subject the system to its most demanding quantitative condition at each sequence, while orthogonal retention time coefficient monitoring isolates chromatographic and instrument-state perturbations. Acceptance criteria (recovery, identification depth, intra-bracket precision) monitor independent failure modes. Revalidation triggers under ICH Q14 lifecycle management are defined prospectively and include sustained SST excursions, database or software version updates altering protein inference by >5%, and extension to matrices outside the validated domain (**Supplementary Method S3; Table S12**).

The validated range was defined in absolute mass units (ng) to decouple performance from product-specific matrix composition. Expression in ppm would bind validation to a specific product modality and, at ppm levels corresponding to purified mAb matrices, the number of identifiable HCPs collapses toward single digits at the lower bound, precluding estimation of population-level variance components or TIs with stable degrees of freedom. An ng-scale range accommodates matrices spanning orders-of-magnitude differences in HCP burden. For purified mAb drug substance, product-specific depletion strategies concentrate the impurity fraction into the validated window; enrichment-related bias should be characterized in auxiliary specificity studies. DIA acquisition eliminates DDA sampling stochasticity and improves per-protein consistency, but two constraints limit immediate regulatory deployment: library-dependent DIA operates within a closed analyte space contingent on prior observation, and no 21 CFR Part 11-compliant software currently supports fully library-free DIA for GMP deployment. Library-free DIA algorithms remove the analyte-space constraint and align with the measurement-system validation framework described here, but the available processing software ecosystem has not matured to support GMP implementation. The TE framework is transferable to DIA once this infrastructure constraint is resolved. The SIL-HCP standard is assumed to approximate aggregate CHO proteome composition based on identical cell-line derivation (**Supplementary Table S13**; **Figure S5**); formal demonstration of peptide-level response-factor equivalence was not performed. Extension to non-CHO expression systems, alternative mAb formats, or alternative processing pipelines requires bridging validation.

Within the defined applicability domain (CHO-derived mAb; SIL-HCP calibrant; 20–80 ng; Evosep One–timsTOF Pro 2 in ddaPASEF; SpectroMine with deterministic parsimony inference), analytical performance satisfied all predefined regulatory criteria. The statistical complexity of the framework, including hierarchical variance decomposition, TI construction, abundance-stratified accuracy profiling, and entrapment-based identification error control, is confined to the validation stage. During routine operation, the procedure reduces to a locked sample preparation protocol, a fixed acquisition method, a version-controlled informatics pipeline, and evaluation of reportable results against predefined acceptance criteria. The SST bracket design provides binary pass/fail decisions at the point of use; the underlying distributional modeling is embedded in the acceptance limits and does not require re-execution.

Untargeted MS-based HCP analysis is unlikely to replace high-throughput immunoassays for routine lot-release screening in the near term. Its principal value lies in providing a platform-level quantitative method with explicit characterization of measurement uncertainty for the impurity population. The framework described here may extend to other analytical domains where analyte identity is determined by computational inference.

## 5. Conclusions

This study presents a prospective, TE-based validation of label-free ddaPASEF proteomics for quantitative host cell protein determination aligned with ICH Q2(R2). Systematic and random error components were estimated independently, propagated into predictive TIs, and evaluated against predefined acceptance limits. All tested levels from 20 to 80 ng total HCP per injection satisfied the ±30% β-expectation containment criterion, defining the validated aggregate analytical range.

Calibration exhibited stable proportional compression (slope ≈ 0.80) with a minor additive offset, attributable to multiplicative contributions from incomplete proteolytic conversion, matrix-driven ionization suppression, Hi3 response-factor mismatch, and deterministic peptide attrition from conservative quality filtering. Precision was dominated by repeatability; between-assay variability was minor relative to the within-assay component and statistically indistinguishable from zero at the available degrees of freedom.

Empirical entrapment analysis demonstrated peptide-level false discovery proportions below 1% at q = 0.01, with geometric attenuation rendering protein-level false identification negligible. Deterministic parsimony inference ensured stability of protein-group definitions for longitudinal comparability. Abundance-stratified evaluation revealed structured calibration heterogeneity concealed in the aggregate metric, with stratum-specific TIs contained within ±35% at all spike levels, defining an abundance-aware LLOQ of 3.6 ppm (P95 = 3.87 ppm). The aggregate total HCP remains the validated reportable measurand.

Robustness assessment under variation of search engine (CCC = 0.998) and MS platform (CCC = 0.980) demonstrated containment within predefined ±30% limits. Platform transfer constitutes a lifecycle change requiring bridging validation. The SST framework, with LLOQ-proximal bracket injections and orthogonal chromatographic monitoring, links validation evidence to continued performance verification under ICH Q14. The validated applicability domain is defined as CHO-derived matrices quantified using SIL-HCP calibration on Evosep One/timsTOF Pro 2 in ddaPASEF mode with version-locked data processing (SpectroMine, deterministic parsimony inference, Hi3 quantification). Extension to alternative expression systems, product formats, MS platforms, or informatics pipelines requires bridging validation.

## Acknowledgments

The authors gratefully acknowledge Nora Zaïm for her contributions to the laboratory work that supported this study.

