## Supplementary for "Prospective ICH Q2(R2)-aligned total-error validation of label-free untargeted proteomics for host cell protein quantification in biotherapeutics"

Corresponding Author: Somar Khalil

### Table of Contents

**Supplementary Method S1** — Locked Analytical Protocol
**Supplementary Method S2** — Deterministic Parsimony Inference Algorithm
**Supplementary Method S3** — System Suitability Testing and Statistical Process Control

**Supplementary Table S1** — Cophenetic correlation coefficients by spike level
**Supplementary Table S2** —Entrapment fraction among accepted peptides at q=0.01
**Supplementary Table S3** — Power analysis for between-assay variance detection
**Supplementary Table S4** — REML variance estimates (Aggregate)
**Supplementary Table S5** — Dual LLOQ summary
**Supplementary Table S6** — Trueness summary for abundance-stratified HCP quantification
**Supplementary Table S7** — Precision summary for abundance-stratified HCP quantification
**Supplementary Table S8** — Accuracy summary for abundance-stratified HCP quantification
**Supplementary Table S9** — REML variance estimates (Stratified)
**Supplementary Table S10** — Missed cleavage distribution
**Supplementary Table S11** — MassPREP response factor dispersion
**Supplementary Table S12** — Revalidation trigger matrix
**Supplementary Table S13** — SIL-HCP physicochemical coverage summary

**Supplementary Figure S1** — REML vs. Method-of-Moments Variance Comparison
**Supplementary Figure S2** — Pilot SST Control Charts
**Supplementary Figure S3** —Distribution of missed cleavages under trypsin/P digestion.
**Supplementary Figure S4** — MassPREP RF Distribution
**Supplementary Figure S5** —Coverage of SIL-HCP detected proteins vs CHO proteome.

**Supplementary Note S1** — Peptide-to-Protein FDP Propagation Bound
**Supplementary Note S2** — Sensitivity Analysis for Hierarchical Design
**Supplementary Note S3** — L4 Anchor Normalization and Absolute Bias Decomposition
**Supplementary Note S4** — Acceptance Limit Derivation
**Supplementary Note S5** — SIL-HCP Calibrant Representativeness Assessment

### Supplementary Method S1 — Locked Analytical Protocol

#### S1.1 Scope and Applicability

This protocol defines the locked analytical procedure applied to all validation assays reported in the accompanying manuscript. The method enables identification and quantification of HCPs by label-free shotgun proteomics using TIMS-ddaPASEF. All procedural parameters, reagent concentrations, instrument settings, and data processing steps were fixed prior to execution of the validation campaign. No protocol modifications were introduced between assays.

The validated applicability domain is CHO-derived matrices quantified using stable isotope-labeled CHO whole-proteome (SIL-HCP) calibration on Evosep One coupled to a timsTOF Pro operated in ddaPASEF mode with version-locked data processing and deterministic parsimony inference.

#### S1.2 Reference Materials and Standards

##### S1.2.1 Matrix

NISTmAb (RM 8671, humanized IgG1κ) was obtained from the National Institute of Standards and Technology (NIST) and served as the product matrix for all validation experiments. The mAb was supplied at a nominal concentration of 10 mg/mL in 12.5 mM L-histidine, 12.5 mM L-histidine hydrochloride monohydrate, pH 6.0.

##### S1.2.2 HCP calibration standard

SILu™CHOP Stable-Isotope Labeled CHO Proteins (Sigma-Aldrich, MSQC12-100UG) were used as the HCP spike material. Each commercial vial (100 µg) was reconstituted in 200 µL of 50 mM Triethylammonium bicarbonate (TEAB) to yield a stock concentration of 0.5 µg/µL. Aliquots were stored at −20 °C until use. The standard comprises *Cricetulus griseus* whole-cell lysate proteins uniformly labeled with $^{13}C_6,^{15}N_2$-lysine and $^{13}C_6,^{15}N_4$-arginine.

##### S1.2.3 Response factor calibration standards

MassPREP Digestion Standards were used for Hi3-based absolute quantification.

| Entry name | SKU | Protein name | Organism | Spike level (fmol/µL) |
|---|---|---|---|---|
| ALBU_BOVIN | 186002329 | P02769 | Bos taurus (Bovine) | 0.40 |
| ADH1_YEAST | 186002328 | P00330 | Saccharomyces cerevisiae (Baker's yeast) | 0.30 |
| PYGM_RABIT | 186002326 | P00489 | Oryctolagus cuniculus (Rabbit) | 0.25 |
| ENO1_YEAST | 186002325 | P00924 | Saccharomyces cerevisiae (Baker's yeast) | 0.37 |

#### S1.3 Spike Design and Sample Preparation

Seven nominal spike levels (L1–L7) were prepared, corresponding to 20, 30, 40, 50, 60, 70, and 80 ng total SIL-HCP protein per injection. At each level, 20 µg of NISTmAb was combined with the appropriate volume of SIL-HCP stock solution. An unspiked control (L0; NISTmAb only) was included to verify absence of SIL-HCP carryover. Three independent preparation replicates were generated at each level within each assay.

##### S1.3.1 Denaturing solution (0.125% RapiGest (w/v) in 50mM TEAB)

Solubilize 1 vial of RapiGest (1 mg) with 1000µl of buffering solution (50 mM TEAB) to give 0.1% (w/v). Store at ambient temperature.

##### S1.3.2 Sample treatment

At each level, 20 µg of NISTmAb was combined with the appropriate volume of SIL-HCP stock solution. An unspiked control (L0; NISTmAb only) was included to verify absence of SIL-HCP carryover. Three independent preparation replicates were generated at each level within each assay.

**Table M1**. Spiking design

| Level | Nominal HCP load (ng) | SIL-HCP spike (0.5µg/µL) vol (µL) | NISTmAb (10µg/µL) vol (µL) | Denaturing solution vol (µL) |
|---|---|---|---|---|
| L1 | 20 | 1.6 | 2 | 190 |
| L2 | 30 | 2.4 | 2 | 190 |
| L3 | 40 | 3.2 | 2 | 190 |
| L4 | 50 | 4.0 | 2 | 190 |
| L5 | 60 | 4.8 | 2 | 190 |
| L6 | 70 | 5.6 | 2 | 190 |
| L7 | 80 | 6.4 | 2 | 190 |

##### S1.3.3 Reduction

5 µL of freshly prepared 500 mM dithiothreitol (DTT) in 50 mM TEAB was added to each sample. Reduction was performed at 50 °C for 40 min with shaking at 450 RPM in a dry bath thermomixer. The final DTT concentration was approximately 12 mM.

##### S1.3.4 Alkylation

5 µL of freshly prepared 1 M iodoacetamide (IAM) in 50 mM TEAB was added to each sample. Alkylation proceeded for 25 min in the dark at room temperature. IAM solutions were discarded after use.

##### S1.3.5 Proteolytic digestion

Trypsin/Lys-C protease mix (MS grade; Thermo Pierce, A41007) was reconstituted in 50 mM TEAB to a concentration of 0.1 µg/µL. For all samples, 5 µL of the protease mixture was added (enzyme-to-substrate ratio 1:40, w/w). Digestion was conducted at 37 °C overnight (~14 h) with shaking at 350 RPM. The reaction was quenched by addition of 1.5 µL of neat TFA, followed by a 20 min hold at 37

°C to hydrolyze acid-labile RapiGest. Samples were centrifuged and supernatants were dried to completeness using a vacuum concentrator.

##### S1.3.6 Peptide desalting

Dried digests were desalted using Pierce Peptide Desalting Spin Columns. Columns were activated with two washes of 300 µL acetonitrile (ACN), centrifuged at 5000 × g for 1 min per wash, followed by two equilibration washes with 300 µL of 0.1% trifluoroacetic acid (TFA) in water. Dried samples were reconstituted in 300 µL of 0.1% TFA, loaded onto the columns, and centrifuged at 3000 × g for 1 min. Three washes with 300 µL of 0.1% TFA were performed. Peptides were eluted with 300 µL of 45% ACN/0.1% TFA, followed by an additional 200 µL elution. Combined eluates were dried to completeness.

##### S1.3.7 Reconstitution and loading

Dried peptides were reconstituted to a concentration of 0.025 µg/µL after including MassPREP mixture spikes. A total of 20 µL (500 ng starting material) was loaded onto conditioned Evotip Pure C18 cartridges per manufacturer instructions. The injection load delivered 5.0, 6.0, 7.5, and 8.0 fmol of PYGM, ADH, ENO, and BSA, respectively.

#### S1.4 Chromatographic and Mass Spectrometric Conditions

##### S1.4.1 Liquid chromatography

**Table M2**. Chromatographic parameters.

| Parameter | Value |
|---|---|
| LC system | Evosep One |
| Method | 30 samples per day (30 SPD) |
| Mobile phase A | 0.1% (v/v) FA in LC-MS water |
| Mobile phase B | 0.1% (v/v) FA in ACN |
| Flow rate | 500 nL/min |
| Column | Evosep Performance EV1137 |
| Column dimensions | 150 mm × 150 µm i.d. |
| Particle size | 1.5 µm |
| Column temperature | 40 °C |

##### S1.4.2 Mass spectrometry

**Table M3**. Mass spectrometric parameters for ddaPASEF acquisition.

| Parameter | Value |
|---|---|
| Instrument | Bruker timsTOF Pro |
| Acquisition mode | ddaPASEF |
| Source | Nano CaptiveSpray ZDV (20 µm emitter) |
| Polarity | Positive |
| Capillary voltage | 1.5 kV |

| | |
|---|---|
| Dry gas flow | 3 L/min |
| Desolvation temperature | 180 °C |
| Funnel 1 RF | 300 Vpp |
| Funnel 2 RF | 200 Vpp |
| Quadrupole ion energy | 5 eV |
| Transfer time | 60 µs |
| PASEF MS/MS scans per cycle | 8 |
| TIMS accumulation/ramp | 130 ms |
| Mass range (m/z) | 100–1350 |
| Ion mobility range ($1/K_0$) | 0.70–1.40 V·s/cm² |
| Target intensity | 25,000 |
| Intensity threshold | 1,000 |
| Active exclusion | Enabled (0.4 min) |
| Cycle time | 1.1 seconds |

#### S1.5 Assay Sequence

Each assay sequence followed a fixed injection sequence: two wash injections, two blank injections, three SST bracket injections (SIL-HCP at L1, 20 ng), two wash injections, two blank injections, followed by analytical samples in triplicate technical injection per biological replicate. The sequence concluded with one wash, one blank, three closing SST bracket injections, two final washes, and one terminal blank. This bracketed design permits assessment of system performance at sequence start and end.

**Table M4**. Assay sequence design

| Sample | Number of injections | Comment |
|---|---|---|
| Wash | 2 | N/A |
| System blank (mobile phase A) | 2 | N/A |
| Thermo, Pierce™ RTC mixture (100 fmol load) | 2 | Second injection used for SST |
| Wash | 2 | N/A |
| System blank | 1 | N/A |
| Control Sample (CS) | 3 | Used for SST (opening bracket) |
| Wash | 2 | N/A |
| System blank | 1 | N/A |
| Sample1 (each sample is followed by 2 Wash and 1 system blank) | 3 | N/A |
| Wash | 2 | N/A |
| System blank | 1 | N/A |
| Sample2 | 3 | N/A |

| | | |
|---|---|---|
| Control Sample (CS) | 3 | Used for SST (closing bracket) |
| Wash | 2 | N/A |
| System blank | 1 | N/A |
| Thermo, Pierce™ RTC mixture (100 fmol load) | 2 | Second injection used for SST (closing bracket) |
| Wash | 2 | N/A |
| System blank | 1 | N/A |

#### S1.6 Database Search and Protein Identification

##### S1.6.1 Search engine and database

Raw data were processed in SpectroMine v5.2 under a fixed parameter set applied uniformly to all assays and spike levels. Trypsin/P was specified with ≤1 missed cleavage. Carbamidomethylation (C) and SIL arginine (+10.008 Da) and lysine (+8.014 Da) were defined as fixed modifications; no variable modifications were allowed. Peptides were restricted to 7–30 amino acids with charge states 2+ to 3+. Mass tolerances were assigned dynamically. Target–decoy competition was used for q-value estimation, and PSMs, peptides, and protein groups were filtered at $q \leq 0.01$. No missing-value imputation was applied. Quantification was based on MS1 extracted ion chromatograms of precursor ions. The protein sequence database comprised: (i) the *Cricetulus griseus* reference proteome (UniProt, taxon ID: 10029); (ii) the NISTmAb heavy and light chain sequences; (iii) and the four MassPREP digest standard sequences (UniProt accessions P02769, P00330, P00489, P00924).

##### S1.6.2 Protein Inference

See Supplementary Method S2 for full algorithmic specification.

#### S1.7 Peptide and Protein Filtering

After protein inference, peptide-level intensities were filtered within each assay and spike level using a predefined QC sequence. Single-hit proteins were excluded. Outlier peptides were identified on $\log_2$-transformed intensities using a modified Z-score41:

$$Z_{\mathrm{i}} = 0.6745\, x_{\mathrm{i}} - xMAD$$

where $x_i$ is the $\log_2$ intensity of peptide i, $\tilde{x}$ the group median, and MAD the median absolute deviation. Peptides with $|Z_i| > 2.8$ were removed. Peptides whose median $\log_2$ intensity differed by more than tenfold from the corresponding protein-level median were excluded. Injection-to-injection loading differences were corrected by TIC scaling within each spike level27. Peptides with technical-triplicate CV > 25% were removed.

#### S1.8 Hi3 Quantification and HCP Mass Estimation

Protein quantification followed the Hi3 approach28, defining protein abundance as the sum of the three most intense peptides per protein at each spike level. Absolute total HCP mass was calculated as:

$$\mathrm{m_{HCP}(ng)} = \left(\frac{\mathrm{I_{HCP}}}{\mathrm{RF}}\right) \mathrm{MW_{HCP}} \times 10^6$$

where $I_{HCP}$ is the summed Hi3 intensity of the HCP ensemble, $RF = \mathrm{median}_k(I_k/n_k)$ is the response factor estimated from MassPREP standards, $I_k$ the Hi3 intensity of standard k, and $n_k$ the injected amount (fmol). $MW_{HCP}$ is the abundance-weighted molecular weight of inferred lead proteins at that level. $I_{HCP}/RF$ yields femtomoles of HCP-equivalent material; multiplication by $MW_{HCP} \times 10^6$ converts to ng.

#### S1.9 Version Control and Traceability

**Table M5**. Software and database version control.

| Component | Version / Specification |
|---|---|
| Acquisition software | Tims Control (Bruker) |
| Primary search engine | SpectroMine 5.2 (Biognosys) |
| Secondary search engine | FragPipe 24.0 / MSFragger 4.4.1 |
| CHO FASTA database | Cricetulus griseus (2801 reviewed entries, UniProt) |
| Protein inference | Deterministic parsimony (Supplementary Method S3) |
| Statistical analysis | Python 3.12 (pinned packages in repository) |
| Random seed | 0 (NumPy default_rng) |
| Bootstrap iterations | 4,000 (hierarchical TI); 10,000 (stratified) |

All analytical parameters were locked prior to execution of the four validation assays. No post hoc optimization of search parameters, filtering thresholds, or quantification logic was performed.

#### S1.10 Reagent Summary

**Table M6**. Reagent inventory.

| Reagent | Supplier | Catalog No. | Storage | Purity/Grade |
|---|---|---|---|---|
| Acetonitrile (LC-MS) | Biosolve | 0001207801BS | Ambient | LC-MS |
| Water (LC-MS) | Biosolve | 0023217823BS | Ambient | LC-MS |
| Formic acid (99%) | Biosolve | 6914131 | 4 °C | ≥99% |
| IAM (no-weigh format) | Thermo | A39271 | Ambient | MS grade |
| DTT (no-weigh format) | Thermo | A39255 | 4 °C | MS grade |
| Trypsin/Lys-C mix | Thermo Pierce | A41007 | −20 °C | MS grade |
| TEAB (1 M) | Thermo | 90114 | 4 °C | ≥99% |
| RapiGest SF | Waters | 186001860 | Ambient | — |
| MassPREP ALBU_BOVIN | Waters | 186002329 | Ambient | — |
| MassPREP ADH1_YEAST | Waters | 186002328 | Ambient | — |
| MassPREP PYGM_RABIT | Waters | 186002326 | Ambient | — |
| ENO1_YEAST | Waters | 186002325 | Ambient | — |
| SILu™CHOP | Sigma | MSQC12-100UG | −20 °C | — |
| Desalting spin columns | Thermo Pierce | 89852 | Ambient | — |
| Evotip Pure C18 | Evosep | — | Ambient | — |

### Supplementary Method S2 — Deterministic Parsimony Inference Algorithm

#### S2.1 Algorithm Specification

**Input:**

- Reference FASTA: *Cricetulus griseus* proteome (UniProt, taxon ID: 10029, 83,419 entries)
- Observed peptide list: Filtered peptide identifications from SpectroMine (post-FDR, post-quality filtering)

**Procedure:**

**Step 1. In silico digestion**. Digest all protein sequences in the reference FASTA using trypsin specificity (cleavage C-terminal to K and R, including when followed by P) with ≤1 missed cleavage. Optional N-terminal methionine excision is applied: for sequences beginning with M, both the intact and Met-excised forms are digested. Peptide length: 7–30 amino acids. Generate a peptide-to-protein mapping $P \to R$, where P is the set of theoretical peptides and R is the set of proteins from which each peptide can be derived.

**Step 2. Evidence matching.** For each observed peptide $p_obs$ in the filtered identification list, retrieve all proteins mapped to $p_{\mathrm{obs}}$ in the theoretical digest. Peptides absent from the theoretical digest (e.g., due to length or specificity constraints) are excluded. This defines the observed peptide set $P_{\mathrm{obs}}$ and the corresponding protein-to-peptide mapping $R \to P_{\mathrm{obs}}$.

**Step 3. Greedy parsimony assignment.**

**(a)** For each protein r, compute the number of currently unassigned peptides it explains:

$$\left|p \in P_{\mathrm{remaining}}: r \in map(p)\right|$$

**(b)** Select the protein $r*$ with the maximum explanatory peptide count. In case of ties, selection follows Python's $max()$ behavior on the iteration order (deterministic for fixed input).

**(c)** Record all peptides explained by $r*$ as a peptide group. Record all proteins capable of explaining any peptide in this group as the candidate protein set for that group.

**(d)** Remove explained peptides from $P_remaining$.

**(e)** Repeat (a)–(d) until $P_{\mathrm{remaining}}$ is empty or no protein explains any remaining peptide.

**Step 4. Unique peptide filtering.** For each peptide group, identify proteins within the candidate set that explain at least one unique peptide (a peptide mapping to exactly one protein in the full peptide-to-protein map). Proteins lacking unique peptide support are removed from the group. If no protein in the group possesses unique peptide support, all candidate proteins are retained.

**Step 5. Lead protein assignment.** Within each filtered protein group, compute the number of group peptides explained by each member protein. The protein explaining the maximum number of peptides is designated the lead accession. The group label is constructed as the semicolon-delimited concatenation of all member accessions (sorted alphabetically).

**Step 6. Output assignment.** Each observed peptide is assigned to its corresponding group label. The lead accession is recorded separately for downstream quantification.

##### S2.2 Shared-Peptide Handling

The greedy selection criterion (Step 3) resolves shared-peptide conflicts by assigning peptides to the protein group that maximizes coverage at each iteration. Shared peptides are assigned exclusively to the first group that claims them; no peptide is assigned to multiple groups. This exclusive assignment ensures that quantification (Hi3) does not double-count peptide intensities. Paralogous gene families generating extensive shared-peptide networks are collapsed into multi-protein groups when no member possesses distinguishing unique peptides. Such groups are reported with all constituent accessions; the lead accession is operationally defined but does not imply biological precedence.

##### S2.3 Protein Group Reporting Conventions

- Each reported protein group is identified by a single lead accession.
- Group membership (all constituent accessions) is recorded in the supplementary protein list.
- The reportable quantity (Hi3 abundance) is attributed to the lead accession.

##### S2.4 Determinism Guarantee

Given identical inputs (FASTA version, peptide evidence list, digestion parameters), the algorithm produces an invariant output.

No stochastic element (random tie-breaking, sampling) is present. Tie resolution in Step 3 follows deterministic iteration order established by the input data structure.

This property provides a stable protein-group denominator for longitudinal and comparability analyses.

Vendor-specific protein grouping heuristics that change between software releases may alter group membership and reported protein counts independent of analytical variation. Deterministic parsimony eliminates this source of non-analytical variability.

##### S2.5 Digestion Parameters

| Parameter | Value |
|---|---|
| Enzyme specificity | Trypsin (cleavage after K/R, proline-permissive) |
| Missed cleavages | ≤1 |
| Minimum peptide length | 7 aa |
| Maximum peptide length | 30 aa |
| N-terminal Met excision | Enabled |

#### S2.6 Code Availability

The repository contains: parsimony_inference.py (core algorithm implementation), requirements.txt (package dependencies), test_data/ (minimal test FASTA and peptide list for verification), and README.md (usage instructions and parameter specification).

### Supplementary Method S3 — System Suitability Testing and Statistical Process Control

#### S3.1 Overview

System suitability testing (SST) was implemented as a two-component framework monitoring quantitative stability and instrument-state stability at each analytical sequence. The design is aligned with ICH Q2(R2) for continued performance verification and ICH Q14 lifecycle principles. Run-level SST metrics were evaluated against predefined specification limits and monitored longitudinally using Phase I Individuals–Moving Range (I-MR) control charts.

#### S3.2 Bracket Design

***Component 1: SIL-HCP QC Bracket***

At the start and end of each analytical sequence, three replicate injections of the SIL-HCP standard at the LLOQ-proximal level (L1, 20 ng total HCP) were acquired. The arithmetic mean of each triplicate defined the bracket-level reportable SST value. Run-level recovery was defined as the mean of the opening and closing bracket values.

***Component 2: Retention Time Calibration (RTC)***

An RTC peptide mixture (100 fmol load) was injected at sequence start and end to monitor chromatographic and MS stability. Twelve peptides from the Pierce™ Peptide Retention Time Calibration Mixture (Cat. No. 88321) were monitored. For each sequence:

- Median retention time deviation (ΔRT, min) was computed relative to the locked validation reference.
- Median MS1 intensity was computed across the monitored peptides.

MS1 performance was evaluated as percent deviation relative to the Phase I baseline median reference.

| Peptide number | Peptide sequence | Mass |
|---|---|---|
| 1 | SSAAPPPPP**R** | 985.5220 |
| 2 | GISNEGQNASI**K** | 1224.6189 |
| 3 | HVLTSIGE**K** | 990.5589 |
| 4 | IGDYAGI**K** | 843.4582 |
| 5 | TASEFDSAIAQD**K** | 1389.6503 |
| 6 | SAAGAFGPELS**R** | 1171.5861 |
| 7 | ELGQSGVDTYLQT**K** | 1545.7766 |
| 8 | GLILVGGYGT**R** | 1114.6374 |
| 9 | SFANQPLEVVYS**K** | 1488.7704 |

| | | |
|---|---|---|
| 10 | LTILEEL**R** | 995.5890 |
| 11 | ELASGLSFPVGF**K** | 1358.7326 |
| 12 | LSSEAPALFQFDL**K** | 1572.8279 |

#### S3.3 Specification Limits

***Quantitative Recovery***

Opening and closing SIL-HCP bracket means were required to fall within 70–130% of the nominal L1 value (±30% TE validation limits). Failure of the opening bracket precluded sequence initiation. Failure of the closing bracket following a passing opening bracket triggered deviation investigation prior to result release.

***RTC Acceptance Criteria***

- Median ΔRT within ±0.50 min
- Median MS1 deviation within ±35% of the Phase I baseline median reference

These criteria isolate chromatographic and sensitivity perturbations independently of digestion and quantification performance.

No identification-depth threshold (e.g., P05) was enforced at run level during pilot SST monitoring.

#### S3.4 Statistical Process Control (Phase I)

***I-MR Charts***

Run-level metrics monitored by I-MR charts:

- SIL-HCP mean recovery (%)
- Protein identification count
- RTC median ΔRT (min)
- RTC median MS1 deviation (%)

Control limits were derived from Phase I baseline data using standard Individuals–Moving Range methodology (moving range span = 2):

- CL = mean of baseline observations
- $\sigma = \bar{MR} / d_2$, with $d_2 = 1.128$
- UCL = CL + 3σ
- LCL = CL − 3σ
- MR UCL = $D_4 \times \bar{MR}$, with $D_4 = 3.267$

Baseline establishment required a minimum of 20–25 independent analytical sequences. During baseline accumulation, control limits were considered provisional. After baseline completion, limits were fixed for continued monitoring until requalification.

Specification limits (e.g., 70–130% recovery; ±0.50 min ΔRT; ±35% MS1 deviation) were evaluated independently of statistical control limits.

No $\bar{X}$/S charts, Western Electric supplementary rules, or process capability (Cpk) calculations were applied in the pilot SST implementation.

##### S3.5 Pilot SST Charts

Pilot SST control charts were populated during the initial operational deployment phase. A minimum of 30 independent sequences was required to finalize Phase I control limits. Finalized limits are reported with the corresponding pilot dataset.

#### Supplementary Table S1 — Cophenetic correlation coefficients by spike level

Preservation of the protein–protein covariance structure was evaluated by two complementary diagnostics applied to Spearman correlation matrices computed on $\log_{10}$-transformed intensities after regression-based removal of spike-level effects. Panel A reports dendrogram fidelity within each spike level. Panel B reports pairwise matrix concordance between levels. The top 1,250 proteins by inter-sample variance were selected per level for Panel A; Panel B was computed on the intersection of 648 proteins common to all five levels to ensure identical dimensionality. Distance matrices were defined as $D = 1 - \rho$. Hierarchical clustering used average linkage. Optimal cluster count was determined by silhouette maximization over k = 2–12 on precomputed distance matrices.

##### Panel A. Cophenetic correlation and cluster structure by spike level

| Spike Level | CCC_coph | Cluster Count | Silhouette Score |
|---|---|---|---|
| L1 | 0.544 | 2 | 0.237 |
| L2 | 0.560 | 2 | 0.251 |
| L4 | 0.600 | 2 | 0.285 |
| L6 | 0.579 | 2 | 0.274 |
| L7 | 0.526 | 2 | 0.240 |

CCC: cophenetic correlation coefficient quantifying the fidelity with which the average-linkage dendrogram preserves pairwise distances in the original correlation matrix. Higher values indicate better dendrogram representation.

##### Panel B. Pairwise matrix concordance (Mantel permutation test)

| $Level_i$ | $Level_j$ | $n_p roteins$ | $Mantel_r$ | $p_{perm}$ |
|---|---|---|---|---|
| L1 | L2 | 648 | 0.206 | < 0.001 |
| L1 | L4 | 648 | 0.144 | < 0.001 |
| L1 | L6 | 648 | 0.085 | < 0.001 |
| L1 | L7 | 648 | 0.062 | < 0.001 |
| L2 | L4 | 648 | 0.251 | < 0.001 |
| L2 | L6 | 648 | 0.137 | < 0.001 |
| L2 | L7 | 648 | 0.083 | < 0.001 |
| L4 | L6 | 648 | 0.272 | < 0.001 |
| L4 | L7 | 648 | 0.168 | < 0.001 |
| L6 | L7 | 648 | 0.298 | < 0.001 |

$Mantel_r$: Pearson correlation between vectorized upper triangles of the pairwise distance matrices $D_i$ and $D_j$. Permutation p-values were obtained by 2,000 node-relabeling permutations; all observed statistics exceeded every permuted value, yielding $p = 1/(2001) < 0.001$.

#### Supplementary Table S2 — Entrapment fraction among accepted peptides at q=0.01 (Wilson 95% CI)

| Entrapment design | q threshold | N_total(q) | N_entrap(q) | Entrapment fraction | Wilson 95% CI |
|---|---|---|---|---|---|
| Shuffled | 0.0100 | 13907 | 50 | 0.0036 | [0.002728, 0.004736] |
| Trimmed | 0.0100 | 13709 | 57 | 0.0042 | [0.003211, 0.005383] |

*Wilson 95% confidence intervals quantify binomial sampling variability of the entrapment fraction at q = 0.01. The narrow interval widths reflect the large number of accepted peptides and correspondingly stable proportion estimates. The observed entrapment fractions (0.36% and 0.42%) are substantially below the nominal 1% threshold.*

### Supplementary Table S3 — Power analysis for between-assay variance detection

| Level | $\sigma_W$(%) | $\sigma_B$,min at 80% power (%) | $df_B$ | $df_W$ | $m_h$ |
|---|---|---|---|---|---|
| L1 | 2.29 | 3.24 | 3 | 8 | 3 |
| L2 | 1.61 | 2.28 | 3 | 8 | 3 |
| L3 | | | 0 | 2 | |
| L4 | 3.09 | 4.37 | 3 | 8 | 3 |
| L5 | | | 0 | 2 | |
| L6 | 1.35 | 1.91 | 3 | 8 | 3 |
| L7 | 3.81 | 5.38 | 3 | 8 | 3 |

*Minimum detectable between-assay SD at 80% power for* $df_B = 3, df_W = 8, \alpha = 0.05$, *harmonic mean* $m_h = 3.0$. *Computed from the noncentral F-test for* $F = MS_B/MS_W$ *with noncentrality* $\lambda = df_B\, m_h(\sigma_B/\sigma_W)^2$. *The required noncentrality parameter "λ₈₀ ≈ 17.96"* $\sigma_B, min = \sigma_W \times \sqrt{(\lambda^{80}/(df_B \times m_h))} = \sigma_W \times \sqrt{(17.96/9)} = \sigma_W \times 1.41$. *L3 and L5 are not estimable.*

### Supplementary Table S4 — REML variance estimates (Aggregate)

| Level | Spike (ng) | MoM $\sigma_W$(%) | MoM $\sigma_B$(%) | REML $\sigma_W$(%) | REML $\sigma_B$ (%) | Note |
|---|---|---|---|---|---|---|
| 1 | 20 | 2.29 | 0.14 | 2.29 | 0.14 | |
| 2 | 30 | 1.61 | 1.67 | 1.61 | 1.67 | |
| 3 | 40 | 2.16 | 0* | 2.16 | N/E | *N/E ($\sigma_B$ only)* |
| 4 | 50 | 3.09 | 0* | 3.01 | 0.4 | |
| 5 | 60 | 0.69 | 0* | 0.69 | N/E | *N/E ($\sigma_B$ only)* |
| 6 | 70 | 1.35 | 1.48 | 1.35 | 1.48 | |
| 7 | 80 | 3.81 | 0* | 3.43 | 0.02 | |

** MoM estimate truncated to zero ($MS_{between} \leq MS_{within}$). N/E = Not estimable (single-assay level). REML estimates are unconstrained and may yield small positive values where MoM truncates to zero.*

### Supplementary Table S5 — Dual LLOQ summary

| Parameter | Aggregate LLOQ | Abundance-Aware LLOQ |
|---|---|---|
| Value | 20 ng total HCP per injection | 3.6 ppm (P95 = 3.87 ppm) |
| Metrological domain | Total HCP burden | Per-stratum mean abundance |
| Units | ng | ppm (ng HCP / mg mAb) |
| Decision rule | 95% β-expectation TI ⊆ ±30% | 95% β-expectation TI ⊆ ±35% (all strata) |
| Analytical question | Method sensitivity for aggregate impurity quantification | Lowest abundance stratum at which population-level quantitative performance is demonstrated |
| ICH Q2(R2) alignment | Section 6 (Quantitation Limit) | Section 6 extended to stratified reporting |
| Interpretation | Reportable result for total burden release/in-process testing | Informational; applicable when abundance-class-level interpretation is required |

*These two limits address distinct analytical questions and are not interchangeable. The aggregate LLOQ defines the minimum reportable total HCP burden. The abundance-aware LLOQ defines the lowest stratum-level mean at which the TE criterion is satisfied.*

#### Supplementary Table S6. Trueness summary for abundance-stratified HCP quantification

| Stratum | Level (L) | Expected abundance (ppm) | Mean observed abundance (ppm) | $Bias_{j,b}$ (%) | Mean recovery (%) | P05-P95 $RE_{ajrb}$ (%) | 95% CI of Recovery (%) |
|---|---|---|---|---|---|---|---|
| Q1 | 1 | 3.10 | 3.62 | 16.83 | 116.83 | [7.40, 24.86] | [112.43, 121.23] |
| Q1 | 2 | 4.65 | 5.15 | 10.79 | 110.79 | [7.25, 15.51] | [108.76, 112.81] |
| Q1 | 3 | 6.20 | 6.86 | 10.69 | 110.69 | [5.34, 16.61] | [95.05, 126.33] |
| Q1 | 4 | 7.75 | 7.75 | 0.00 | 100.00 | [-9.31, 9.40] | [95.85, 104.15] |
| Q1 | 5 | 9.30 | 10.27 | 10.39 | 110.39 | [8.82, 11.57] | [106.41, 114.37] |
| Q1 | 6 | 10.85 | 11.59 | 6.86 | 106.86 | [-2.09, 16.90] | [102.60, 111.12] |
| Q1 | 7 | 12.40 | 13.83 | 11.57 | 111.57 | [1.77, 22.63] | [107.06, 116.07] |
| Q2 | 1 | 7.27 | 8.47 | 16.53 | 116.53 | [8.41, 24.18] | [112.81, 120.25] |
| Q2 | 2 | 10.90 | 11.31 | 3.75 | 103.75 | [-0.41, 9.04] | [101.55, 105.95] |
| Q2 | 3 | 14.53 | 14.53 | -0.02 | 99.98 | [-1.69, 2.24] | [94.27, 105.69] |
| Q2 | 4 | 18.17 | 18.17 | 0.00 | 100.00 | [-7.10, 6.65] | [96.70, 103.30] |
| Q2 | 5 | 21.80 | 23.14 | 6.17 | 106.17 | [4.73, 7.06] | [102.53, 109.81] |
| Q2 | 6 | 25.43 | 25.81 | 1.49 | 101.49 | [-5.73, 6.89] | [98.64, 104.35] |
| Q2 | 7 | 29.07 | 30.13 | 3.68 | 103.68 | [-5.09, 15.09] | [99.14, 108.22] |
| Q3 | 1 | 16.48 | 18.53 | 12.44 | 112.44 | [9.46, 15.54] | [111.03, 113.85] |
| Q3 | 2 | 24.72 | 26.29 | 6.36 | 106.36 | [1.65, 9.42] | [104.38, 108.34] |
| Q3 | 3 | 32.95 | 33.50 | 1.66 | 101.66 | [-2.21, 5.98] | [90.28, 113.04] |
| Q3 | 4 | 41.19 | 41.19 | 0.00 | 100.00 | [-7.55, 6.24] | [96.71, 103.29] |
| Q3 | 5 | 49.43 | 48.45 | -1.98 | 98.02 | [-2.76, -1.39] | [96.03, 100.01] |
| Q3 | 6 | 57.67 | 55.22 | -4.24 | 95.76 | [-8.01, -1.30] | [94.20, 97.31] |
| Q3 | 7 | 65.91 | 63.20 | -4.11 | 95.89 | [-10.22, 2.52] | [93.26, 98.53] |
| Q4 | 1 | 65.25 | 80.30 | 23.07 | 123.07 | [19.63, 26.36] | [121.57, 124.57] |
| Q4 | 2 | 97.87 | 110.65 | 13.06 | 113.06 | [9.49, 16.13] | [111.56, 114.56] |
| Q4 | 3 | 130.49 | 135.98 | 4.21 | 104.21 | [2.70, 6.12] | [99.32, 109.10] |
| Q4 | 4 | 163.11 | 163.11 | 0.00 | 100.00 | [-4.43, 5.05] | [97.88, 102.12] |
| Q4 | 5 | 195.74 | 187.97 | -3.97 | 96.03 | [-4.73, -3.05] | [93.66, 98.40] |
| Q4 | 6 | 228.36 | 209.16 | -8.41 | 91.59 | [-10.64, -6.04] | [90.60, 92.58] |
| Q4 | 7 | 260.98 | 232.57 | -10.89 | 89.11 | [-14.56, -5.91] | [87.22, 91.00] |

#### Supplementary Table S7. Precision summary for abundance-stratified HCP quantification

| Stratum | Level (L) | Expected abundance (ppm) | Within-assay SD (%) | Between-assay SD (%) | Total SD (%) | Within-assay df | Between-assay df |
|---|---|---|---|---|---|---|---|
| Q1 | 1 | 3.10 | 8.06 | 0.00 | 8.06 | 8 | 3 |
| Q1 | 2 | 4.65 | 2.29 | 2.45 | 3.35 | 8 | 3 |
| Q1 | 3 | 6.20 | 6.30 | 0.00 | 6.30 | 2 | 0 |
| Q1 | 4 | 7.75 | 5.86 | 3.20 | 6.67 | 8 | 3 |
| Q1 | 5 | 9.30 | 1.60 | 0.00 | 1.60 | 2 | 0 |
| Q1 | 6 | 10.85 | 5.86 | 3.59 | 6.88 | 8 | 3 |
| Q1 | 7 | 12.40 | 5.42 | 5.07 | 7.42 | 8 | 3 |
| Q2 | 1 | 7.27 | 5.03 | 3.32 | 6.02 | 8 | 3 |
| Q2 | 2 | 10.90 | 2.94 | 2.02 | 3.57 | 8 | 3 |
| Q2 | 3 | 14.53 | 2.30 | 0.00 | 2.30 | 2 | 0 |
| Q2 | 4 | 18.17 | 4.31 | 3.20 | 5.37 | 8 | 3 |
| Q2 | 5 | 21.80 | 1.46 | 0.00 | 1.46 | 2 | 0 |
| Q2 | 6 | 25.43 | 2.42 | 4.17 | 4.83 | 8 | 3 |
| Q2 | 7 | 29.07 | 7.92 | 0.00 | 7.92 | 8 | 3 |
| Q3 | 1 | 16.48 | 2.34 | 0.00 | 2.34 | 8 | 3 |
| Q3 | 2 | 24.72 | 2.00 | 2.64 | 3.31 | 8 | 3 |
| Q3 | 3 | 32.95 | 4.58 | 0.00 | 4.58 | 2 | 0 |
| Q3 | 4 | 41.19 | 4.14 | 3.42 | 5.37 | 8 | 3 |
| Q3 | 5 | 49.43 | 0.80 | 0.00 | 0.80 | 2 | 0 |
| Q3 | 6 | 57.67 | 1.64 | 2.01 | 2.60 | 8 | 3 |
| Q3 | 7 | 65.91 | 4.60 | 0.00 | 4.60 | 8 | 3 |
| Q4 | 1 | 65.25 | 2.46 | 0.00 | 2.46 | 8 | 3 |
| Q4 | 2 | 97.87 | 1.83 | 1.63 | 2.46 | 8 | 3 |
| Q4 | 3 | 130.49 | 1.97 | 0.00 | 1.97 | 2 | 0 |
| Q4 | 4 | 163.11 | 2.91 | 1.79 | 3.42 | 8 | 3 |
| Q4 | 5 | 195.74 | 0.96 | 0.00 | 0.96 | 2 | 0 |
| Q4 | 6 | 228.36 | 1.34 | 0.87 | 1.60 | 8 | 3 |
| Q4 | 7 | 260.98 | 3.08 | 0.00 | 3.08 | 8 | 3 |

#### Supplementary Table S8. Accuracy summary for abundance-stratified HCP quantification

| Stratum | Level (L) | Expected abundance (ppm) | Mean observed abundance (ppm) | $Bias_{j,b}$ (%) | Pred. total SD (%) | Satterthwaite df | 95 β-expectation TI (%) |
|---|---|---|---|---|---|---|---|
| Q1 | 1 | 3.10 | 3.62 | 16.83 | 5.53 | 8.00 | [6.0 , 27.7] |
| Q1 | 2 | 4.65 | 5.15 | 10.79 | 5.73 | 5.87 | [-0.4 , 22.0] |
| Q1 | 3 | 6.20 | 6.86 | 10.69 | 5.87 | 3.00 | [-0.8 , 22.2] |
| Q1 | 4 | 7.75 | 7.75 | 0.00 | 5.99 | 8.94 | [-11.7 , 11.7] |
| Q1 | 5 | 9.30 | 10.27 | 10.39 | 6.09 | 3.00 | [-1.5 , 22.3] |
| Q1 | 6 | 10.85 | 11.59 | 6.86 | 6.17 | 8.48 | [-5.2 , 19.0] |
| Q1 | 7 | 12.40 | 13.83 | 11.57 | 6.24 | 6.48 | [-0.7 , 23.8] |
| Q2 | 1 | 7.27 | 8.47 | 16.53 | 4.45 | 8.15 | [7.8 , 25.3] |
| Q2 | 2 | 10.90 | 11.31 | 3.75 | 4.71 | 7.97 | [-5.5 , 13.0] |
| Q2 | 3 | 14.53 | 14.53 | -0.02 | 4.91 | 3.00 | [-9.6 , 9.6] |
| Q2 | 4 | 18.17 | 18.17 | 0.00 | 5.06 | 7.60 | [-9.9 , 9.9] |
| Q2 | 5 | 21.80 | 23.14 | 6.17 | 5.20 | 3.00 | [-4.0 , 16.4] |
| Q2 | 6 | 25.43 | 25.81 | 1.49 | 5.31 | 4.27 | [-8.9 , 11.9] |
| Q2 | 7 | 29.07 | 30.13 | 3.68 | 5.41 | 8.00 | [-6.9 , 14.3] |
| Q3 | 1 | 16.48 | 18.53 | 12.44 | 2.74 | 8.00 | [7.1 , 17.8] |
| Q3 | 2 | 24.72 | 26.29 | 6.36 | 3.02 | 5.03 | [0.4 , 12.3] |
| Q3 | 3 | 32.95 | 33.50 | 1.66 | 3.24 | 3.00 | [-4.7 , 8.0] |
| Q3 | 4 | 41.19 | 41.19 | 0.00 | 3.42 | 7.09 | [-6.7 , 6.7] |
| Q3 | 5 | 49.43 | 48.45 | -1.98 | 3.58 | 3.00 | [-9.0 , 5.0] |
| Q3 | 6 | 57.67 | 55.22 | -4.24 | 3.71 | 5.31 | [-11.5 , 3.0] |
| Q3 | 7 | 65.91 | 63.20 | -4.11 | 3.83 | 8.00 | [-11.6 , 3.4] |
| Q4 | 1 | 65.25 | 80.30 | 23.07 | 2.52 | 8.00 | [18.1 , 28.0] |
| Q4 | 2 | 97.87 | 110.65 | 13.06 | 2.47 | 6.72 | [8.2 , 17.9] |
| Q4 | 3 | 130.49 | 135.98 | 4.21 | 2.43 | 3.00 | [-0.6 , 9.0] |
| Q4 | 4 | 163.11 | 163.11 | 0.00 | 2.41 | 8.46 | [-4.7 , 4.7] |
| Q4 | 5 | 195.74 | 187.97 | -3.97 | 2.38 | 3.00 | [-8.6 , 0.7] |
| Q4 | 6 | 228.36 | 209.16 | -8.41 | 2.37 | 8.23 | [-13.0 , -3.8] |
| Q4 | 7 | 260.98 | 232.57 | -10.89 | 2.35 | 8.00 | [-15.5 , -6.3] |

#### Supplementary Table S9 — REML variance estimates (Stratified)

| Stratum | Level | MoM $\sigma_W$(%) | MoM $\sigma_B$(%) | REML $\sigma_W$(%) | REML $\sigma_B$ (%) | Note |
|---|---|---|---|---|---|---|
| Q1 | 1 | 8.06 | 0 | 6.92 | 0.06 | |
| Q1 | 2 | 2.29 | 2.45 | 2.29 | 2.45 | |
| Q1 | 3 | 6.30 | 0 | 6.30 | N/E | N/E ($\sigma_B$ only) |
| Q1 | 4 | 5.86 | 3.2 | 6.53 | 0 | |
| Q1 | 5 | 1.60 | 0 | 1.6 | N/E | N/E ($\sigma_B$ only) |
| Q1 | 6 | 5.86 | 3.59 | 5.86 | 3.59 | |
| Q1 | 7 | 5.42 | 5.07 | 5.42 | 5.07 | |
| Q2 | 1 | 5.03 | 3.32 | 5.03 | 3.32 | |
| Q2 | 2 | 2.94 | 2.02 | 2.94 | 2.02 | |
| Q2 | 3 | 2.30 | 0 | 2.30 | N/E | N/E ($\sigma_B$ only) |
| Q2 | 4 | 4.31 | 3.20 | 4.31 | 3.20 | |
| Q2 | 5 | 1.46 | 0 | 1.46 | N/E | N/E ($\sigma_B$ only) |
| Q2 | 6 | 2.42 | 4.17 | 2.42 | 4.17 | |
| Q2 | 7 | 7.92 | 0 | 8.11 | 0 | |
| Q3 | 1 | 2.34 | 0 | 2.22 | 0.03 | |
| Q3 | 2 | 2.00 | 2.64 | 2.00 | 2.64 | |
| Q3 | 3 | 4.58 | 0 | 4.58 | N/E | N/E ($\sigma_B$ only) |
| Q3 | 4 | 4.14 | 3.42 | 4.14 | 3.42 | |
| Q3 | 5 | 0.80 | 0 | 0.80 | N/E | N/E ($\sigma_B$ only) |
| Q3 | 6 | 1.64 | 2.01 | 1.64 | 2.01 | |
| Q3 | 7 | 4.60 | 0 | 4.09 | 1.26 | |
| Q4 | 1 | 2.46 | 0 | 2.37 | 0 | |
| Q4 | 2 | 1.83 | 1.63 | 1.83 | 1.63 | |
| Q4 | 3 | 1.97 | 0 | 1.97 | N/E | N/E ($\sigma_B$ only) |
| Q4 | 4 | 2.91 | 1.79 | 2.91 | 1.79 | |
| Q4 | 5 | 0.96 | 0 | 0.96 | N/E | N/E ($\sigma_B$ only) |
| Q4 | 6 | 1.34 | 0.87 | 1.34 | 0.87 | |
| Q4 | 7 | 3.08 | 0 | 2.98 | 0 | |

** MoM estimate truncated to zero ($MS_{between} \leq MS_{within}$). N/E = Not estimable (single-assay level). REML estimates are unconstrained and may yield small positive values where MoM truncates to zero.*

### Supplementary Table S10 — Missed cleavage distribution

| Missed cleavages | Number of peptides | Percentage (%) |
|---|---|---|
| 0 | 19096 | 95.81 |
| 1 | 835 | 4.19 |

*With missed-cleavage fraction <15%, residual bias likely originates from ionization suppression and RF mismatch.*

#### Supplementary Table S11 — MassPREP response factor dispersion

| Assay | Standard protein | $RF_{L1}$ | $RF_{L2}$ | $RF_{L3}$ | $RF_{L4}$ | $RF_{L5}$ | $RF_{L6}$ | $RF_{L7}$ |
|---|---|---|---|---|---|---|---|---|
| 1 | ADH1_YEAST | 2.60E+03 | 2.45E+03 | 3.03E+03 | 2.36E+03 | 2.43E+03 | 2.77E+03 | 2.59E+03 |
| 2 | | 2.64E+03 | 2.95E+03 | | 2.80E+03 | | 2.83E+03 | 2.71E+03 |
| 3 | | 3.17E+03 | 2.69E+03 | | 2.36E+03 | | 2.73E+03 | 2.66E+03 |
| 4 | | 3.02E+03 | 3.00E+03 | | 2.58E+03 | | 3.09E+03 | 2.61E+03 |
| 1 | ALBU_BOVIN | 2.44E+03 | 2.47E+03 | 2.47E+03 | 2.34E+03 | 2.30E+03 | 2.35E+03 | 2.22E+03 |
| 2 | | 2.42E+03 | 2.39E+03 | | 2.27E+03 | | 2.24E+03 | 2.17E+03 |
| 3 | | 2.47E+03 | 2.38E+03 | | 2.25E+03 | | 2.34E+03 | 2.32E+03 |
| 4 | | 2.44E+03 | 2.43E+03 | | 2.49E+03 | | 2.58E+03 | 2.50E+03 |
| 1 | ENO1_YEAST | 2.69E+03 | 2.53E+03 | 2.52E+03 | 2.41E+03 | 2.33E+03 | 2.29E+03 | 2.23E+03 |
| 2 | | 2.83E+03 | 2.80E+03 | | 2.59E+03 | | 2.63E+03 | 2.48E+03 |
| 3 | | 2.99E+03 | 2.86E+03 | | 2.70E+03 | | 2.71E+03 | 2.39E+03 |
| 4 | | 2.96E+03 | 2.93E+03 | | 2.55E+03 | | 2.69E+03 | 2.78E+03 |
| 1 | PYGM_RABIT | 2.69E+03 | 2.39E+03 | 2.40E+03 | 2.44E+03 | 2.43E+03 | 2.35E+03 | 2.16E+03 |
| 2 | | 2.87E+03 | 2.72E+03 | | 2.48E+03 | | 2.31E+03 | 2.44E+03 |
| 3 | | 2.79E+03 | 2.88E+03 | | 2.65E+03 | | 2.31E+03 | 2.46E+03 |
| 4 | | 2.87E+03 | 2.85E+03 | | 2.80E+03 | | 2.30E+03 | 2.10E+03 |
| **Median RF** | | 2.74E+03 | 2.71E+03 | 2.50E+03 | 2.48E+03 | 2.38E+03 | 2.46E+03 | 2.45E+03 |
| **Mean RF** | | 2.74E+03 | 2.67E+03 | 2.60E+03 | 2.50E+03 | 2.37E+03 | 2.53E+03 | 2.43E+03 |
| **CV RF (%)** | | 8.5 | 8.7 | 11.0 | 7.0 | 2.8 | 10.0 | 8.6 |

*With CV < 20%, the moderate dispersion supports the use of the median RF as representative.*

### Supplementary Table S12 — Revalidation trigger matrix

| Trigger Event | Trigger Criterion | Action |
|---|---|---|
| SST trend excursion | 3σ breach on recovery or protein count for 2 consecutive sequences | Investigate; if systematic, partial revalidation (precision + trueness at L1) |
| Column lot change | RT shift exceeding RTC acceptance criterion after equilibration | SST verification run; if fail, partial revalidation |
| Software version update | Protein inference output changes by >5% of reported protein groups | Full reprocessing of validation dataset; assess TI containment |
| Major instrument maintenance | Source replacement, detector recalibration, or firmware update producing SST bracket failure | SST bracket + L1/L4/L7 verification; if fail, partial revalidation |
| New product matrix | Extension to non-CHO host system, alternative mAb format, vaccine matrix, or different MS platform | Full bridging validation with empirical demonstration of comparable variance structure |
| Database update | Reference FASTA version change altering >2% of protein groups | Reprocess validation data; if TIs change, assess containment |
| Cpk trend* | Cpk trending below 1.33 for recovery or protein count | Investigate root cause; preventive maintenance |

*Defined under ICH Q14 analytical procedure lifecycle management. * applies to Phase II monitoring only.*

### Supplementary Table S13 — SIL-HCP physicochemical coverage summary

| Property | SIL-HCP (detected) Median [IQR] [95% CI] | CHO proteome Median [IQR] [95% CI] | Δmedian(det−CHO) [95% CI] |
|---|---|---|---|
| MW (kDa) | 46.01 [28.12, 77.34] | 44.43 [24.08, 79.04] | 1.58 [-0.06, 3.04] |
| pI (theoretical) | 6.51 [5.17, 7.89] | 7.34 [5.98, 8.75] | -0.83 [-0.91, -0.76] |
| GRAVY index | -0.402 [-0.620, -0.225] | -0.376 [-0.618, -0.136] | -0.025 [-0.039, -0.012] |

### Supplementary Figures

#### Supplementary Figure S1 — REML vs. Method-of-Moments Variance Comparison

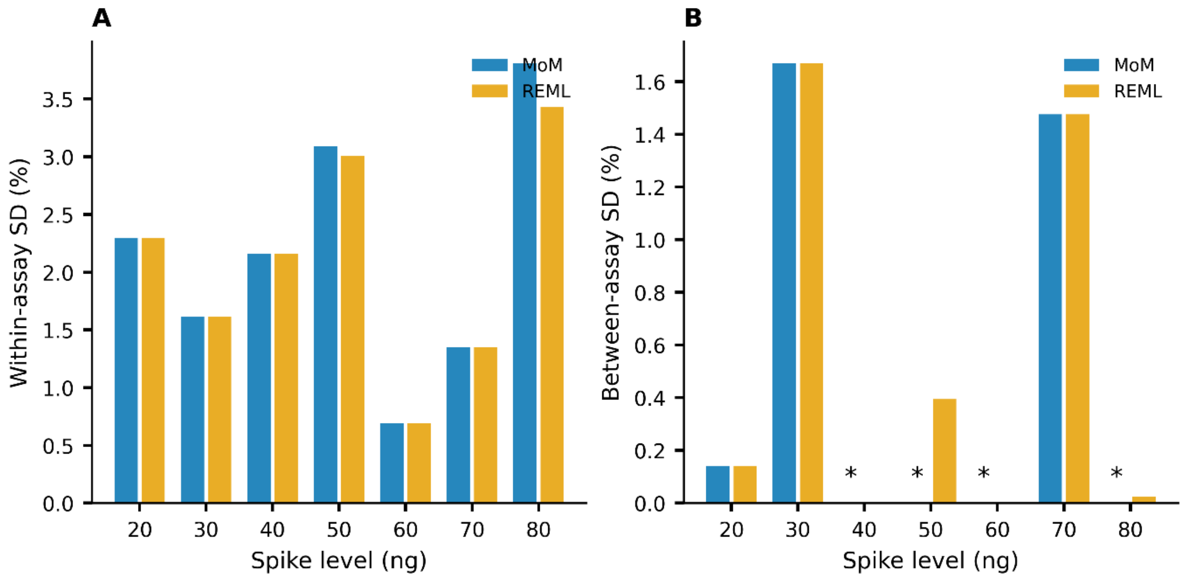


**A**) Within-assay precision expressed as CV% for MoM and REML estimators across spike levels (20–80 ng). (**B**) Between-assay SD for the same estimators across assays at each spike level. Asterisks denote levels where between-assay variance was not estimable or approached zero under the variance-component model.

**Supplementary Figure S2 — Moving Range (MR) charts for pilot SST metrics.**

Pilot SST control charts were populated during the initial operational deployment phase. A baseline of 30 independent analytical sequences was used to establish Phase I control limits. Following baseline establishment, control limits were fixed for ongoing Phase II monitoring.

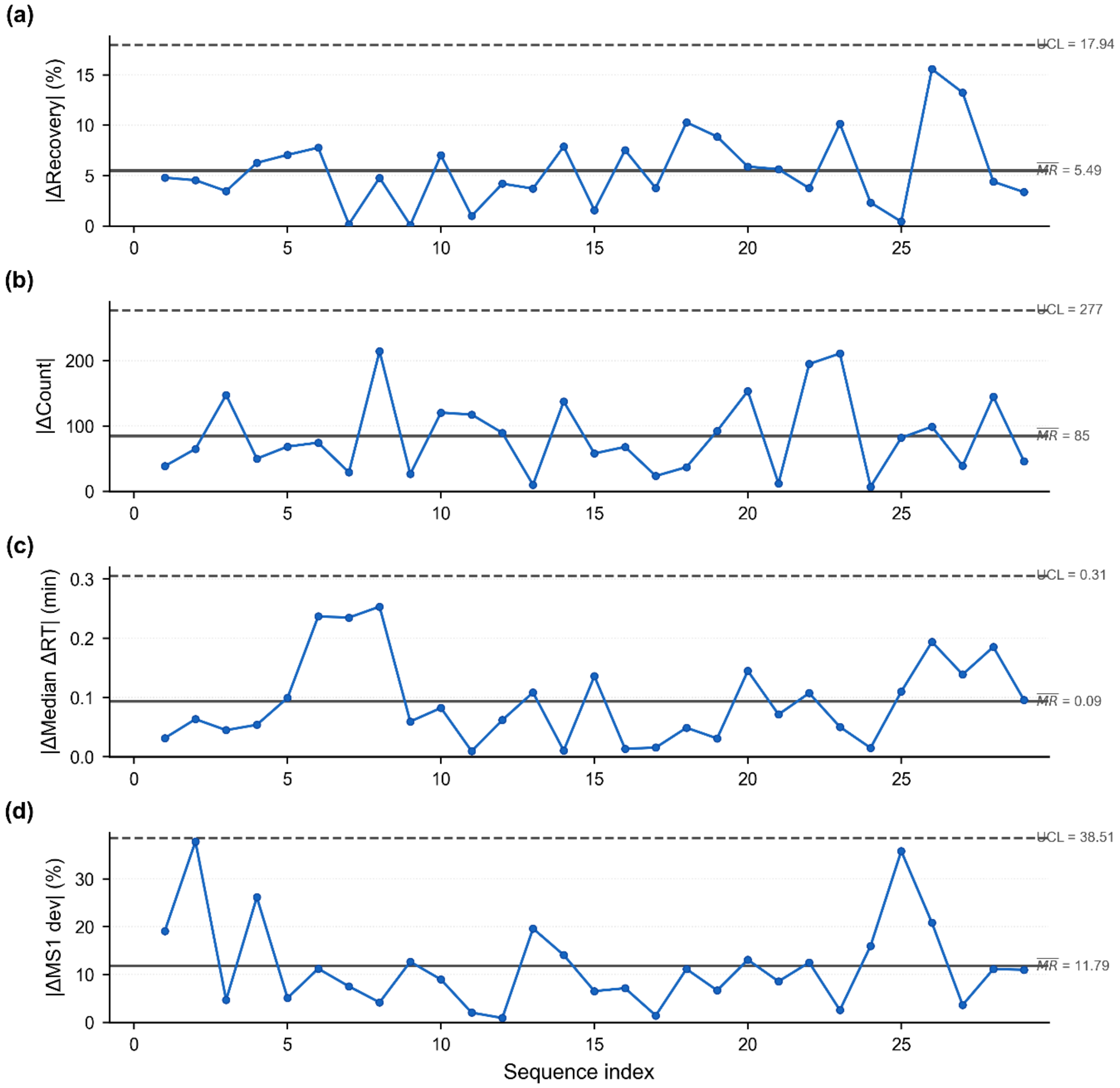


***Phase I moving range** (**a**) SIL-HCP recovery (%), (**b**) protein identification count, (**c**) RTC median ΔRT (min), and (**d**) RTC MS1 deviation (%). The solid line represents the moving range for each consecutive sequence pair; the center line ($\overline{MR}$) and upper control limit (UCL_MR = $D_4 \times \overline{MR}$, $D_4 = 3.267$) are estimated from baseline data using standard I-MR methodology ($d_2$ = 1.128).*

**Supplementary Figure S3 —Distribution of missed cleavages under trypsin/P digestion.**

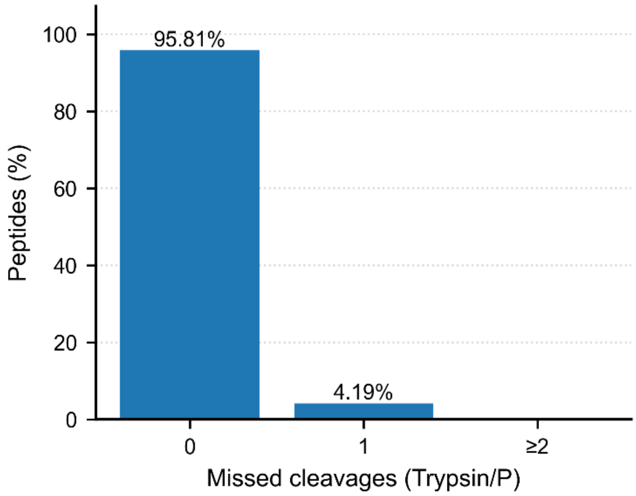


Percentage of identified peptides stratified by the number of internal missed cleavage sites (trypsin/P specificity). Fully tryptic peptides (0 missed cleavages) predominate, with a minor fraction containing a single missed cleavage.

#### Supplementary Figure S4 — MassPREP RF Distribution

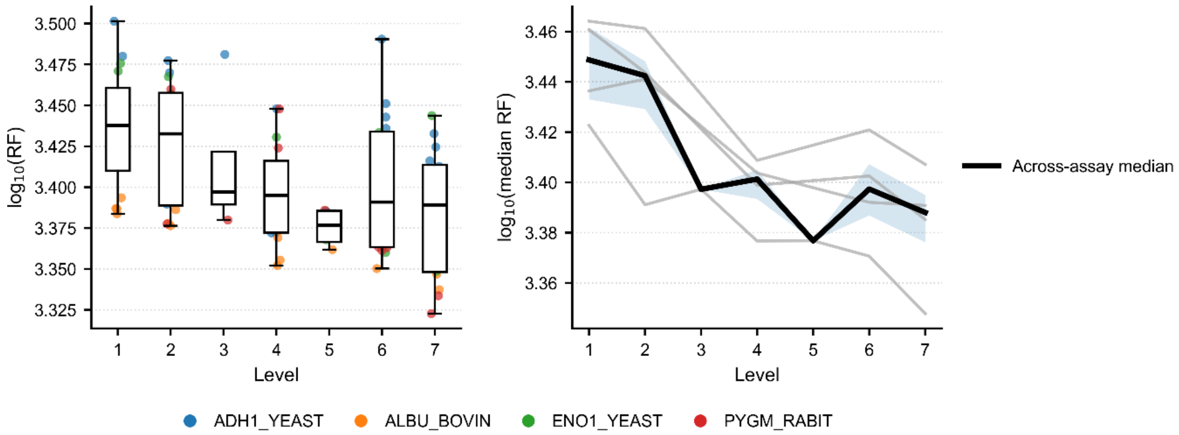


**Left panel:** Distribution of $\log_{10}$-transformed response factors (RF) for each spike level (1–7), pooled across assays. Boxes denote median and interquartile range; whiskers represent the range excluding outliers. Individual points are colored by protein identity. **Right panel:** Median $\log_{10}$(RF) per level for each assay (thin grey lines). The bold black line represents the across-assay median of assay-specific medians at each level, with shaded band indicating the interquartile range across assays.

**Supplementary Figure S5 —Coverage of SIL-HCP detected proteins vs CHO proteome.**

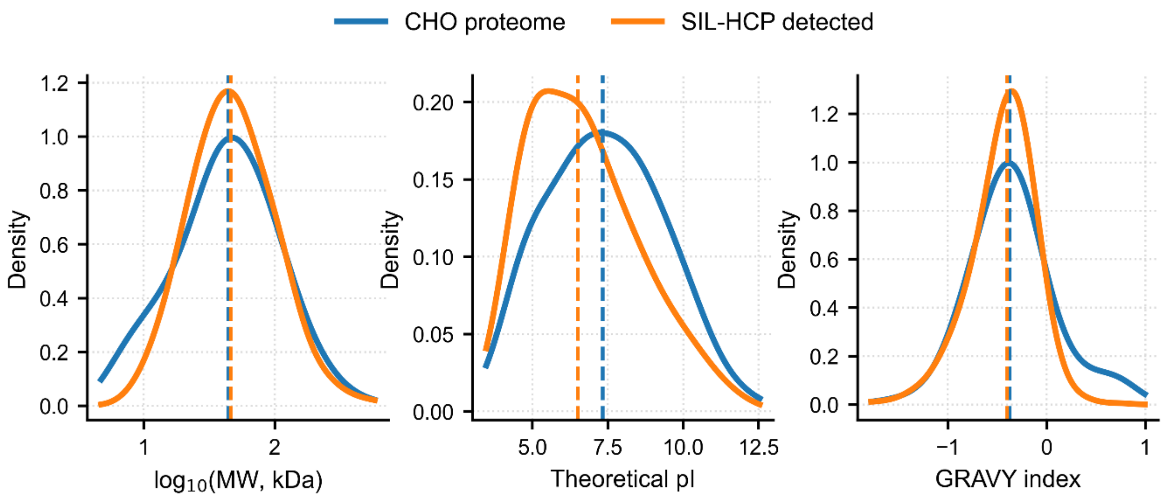


Kernel density estimates of $\log_{10}$-transformed MW (kDa), theoretical isoelectric point (pI), and GRAVY index are shown for the full CHO proteome and the SIL-HCP detected subset. Dashed vertical lines denote distribution medians.

#### Supplementary Note S1 — Peptide-to-Protein FDP Propagation Bound

The entrapment FDP of 0.7–0.9% is estimated at the peptide level. Propagation to the protein level is attenuated by the Hi3 aggregation requirement, which selects the three most intense peptides per protein group for quantification.

For a false protein identification to contribute quantitative error, at least three false peptides must: (a) pass the $q = 0.01$ threshold, (b) map to the same protein group under parsimony inference, and (c) rank among the top three by intensity within that group.

If peptide-level false identifications are modeled as independent Bernoulli events with probability $p = \mathrm{FDP} \approx 0.009$, the probability that $k \geq 3$ concordant false peptides are assigned to the same protein group scales as:

$$\mathrm{P(falseprotein)} \approx p^3 \approx (0.009)^3 \approx 7.3 \times 10^{-7}$$

This is an upper bound because it ignores the additional requirement that all three false peptides must map to the same parsimony group (geometric dilution over the protein space). The parsimony algorithm distributes false peptides over many groups, counteracting any concentration effect from shared-peptide connectivity. The expected protein-level false identification rate is on the order of $10^{-7}$ to $10^{-5}$, negligible relative to other uncertainty components (total SD 0.7–3.8%).

#### Supplementary Note S2 — Sensitivity Analysis for Hierarchical Design

##### S2.1 Context

The four-assay hierarchical design provides df = 3 for between-assay variance estimation. Zero between-assay variance estimates may reflect genuinely negligible variability or limited resolving power.

##### S2.2 Minimum detectable between-assay SD

Under a balanced one-way random-effects model, the F-statistic $MS_b etween/MS_w ithin$ follows F(3, 8) under $H_0$ ($\sigma_B = 0$) and the noncentral F distribution under $H_1$ with noncentrality $\lambda = df_B \cdot m_h \cdot (\sigma_B/\sigma_W)^2$. At α = 0.05 and 80% power, the required noncentrality parameter $\lambda^{80} \approx 17.96$ (at $df^1 = 3$, $df^2 = 8$). Therefore:

$$\sigma_B, min = \sigma_W \times \sqrt{\left(\lambda^{80}/(df_B \times m_h)\right)} = \sigma_W \times \sqrt{(17.96/9)} = \sigma_W \times 1.41$$

Level-specific minimum detectable between-assay SDs are reported in **Supplementary Table S3**.

##### S2.3 Interpretation

Between-assay variance components below the $\sigma_B, min$ thresholds are statistically indistinguishable from zero under the present design. The zero MoM estimates are consistent with $\sigma_B$ values in the range [0, $\sigma_B, min$]. Even if between-assay SD were at the detection threshold (e.g., $\sigma_B \approx 3.2$ at L1), the total SD would be $\sqrt{(2.29^2 + 3.24^2)} = 3.97$, and the β-expectation TI would remain within ±30%.

An expanded design (8–10 assays) would improve resolution but was not justified given the low total SD and margin to acceptance limits.

#### Supplementary Note S3 — L4 Anchor Normalization and Absolute Bias Decomposition

The stratified total-error analysis normalizes observed protein abundances to L4. This normalization forces zero bias at the anchor level by construction. In the aggregate analysis, L4 carries approximately −16.7% bias relative to the nominal gravimetric spike value (Table 4: 50 ng nominal, 41.67 ng observed, recovery 83.34%).

The stratified bias at any spike level j within stratum b is interpretable as deviation from L4-specific systematic error, not from absolute ground truth. The relationship is:

$$\mathrm{Bias}_{\mathrm{a}}\mathrm{bsolute}(\mathrm{j},\mathrm{b}) = \mathrm{Bias}_{\mathrm{L}}4_{\mathrm{a}}\mathrm{ggregate} + \mathrm{Bias}_{\mathrm{s}}\mathrm{tratified}(\mathrm{j},\mathrm{b})$$

**Example 1 (Q1, L1):** $\mathrm{Bias}_{\mathrm{s}}\mathrm{tratified} = +16.83$. $\mathrm{Bias}_{\mathrm{a}}\mathrm{bsolute} = -16.7$. The positive stratified bias does not indicate over-recovery in absolute terms; Q1 proteins at L1 recover approximately 17 percentage points more than the L4 anchor. Because the L4 anchor itself under-recovers by ~17%, absolute recovery of Q1 at L1 is near 100%.

**Example 2 (Q4, L7):** $\mathrm{Bias}_{\mathrm{s}}\mathrm{tratified} = -10.89$. $\mathrm{Bias}_{\mathrm{a}}\mathrm{bsolute} = -16.7$. Q4 at L7 shows the most negative absolute bias, reflecting additive compression from both the aggregate calibration offset and the high-abundance slope deficit.

The stratified TE decision criterion (±35%) is applied to the stratified bias, which is the operationally relevant quantity for within-method comparisons. Absolute recovery is recoverable from the aggregate calibration function.

#### Supplementary Note S4 — Acceptance Limit Derivation

The ±30% acceptance limit for aggregate total-error was derived from the immunoassay-based HCP performance envelope established in USP ⟨1132⟩. The ±30% aggregate acceptance limit was derived from the USP ⟨1132⟩ immunoassay performance envelope and to remain compatible with SFSTP β-expectation guidance.

The ±35% acceptance limit for abundance-stratified total-error accounts for the additional variance introduced by: (i) population-level bootstrap averaging within fixed abundance strata containing tens to hundreds of proteins; (ii) stratum-to-stratum heterogeneity in calibration slope; and (iii) the reduced protein count in lower-abundance strata (Q1), which inflates the contribution of individual protein variability to the stratum mean. The 5 percentage-point widening relative to the aggregate limit accommodates these structural sources of additional dispersion while maintaining regulatory stringency.

### Supplementary Note S5 — SIL-HCP Calibrant Representativeness Assessment

#### S5.1 Compositional basis

The SIL-HCP whole-proteome standard is derived from CHO-K1 cells cultured in heavy-labeled media. The endogenous HCP population in the NISTmAb matrix is derived from a potentially different CHO lineage. Compositional mismatch does not invalidate aggregate-level calibration because: (i) Hi3 quantification aggregates peptide-level signals over hundreds of proteins, attenuating individual protein-level abundance differences; (ii) the calibration model captures the net systematic effect inclusive of any compositional mismatch; (iii) product-specific calibrant matching is desirable for single-protein quantification but not required for aggregate burden estimation under the total-error framework.

#### S5.2 Physicochemical comparison

See Supplementary **Table S13** and Supplementary **Figure S5**. Physicochemical distribution comparisons are necessary but insufficient; the actual validation requirement is response-factor equivalence at the peptide level. This equivalence is not demonstrated empirically in the present work.

#### S5.3 Risk-based justification

Under ICH Q14, calibrant representativeness is evaluated as a risk factor. For aggregate HCP quantification: the risk of compositional mismatch biasing the summed HCP estimate is captured in the WLS calibration model and bounded within ±30% TE. The validation demonstrates that the net effect of all systematic contributions is within acceptance limits. Aggregate quantification is robust to individual protein-level deviations by the central limit theorem argument applied to the Hi3 sum.

#### Code Deposition

The complete statistical analysis pipeline (Python 3.12) is deposited at: **10.5281/zenodo.18826281**
